# Purification protocol of hypertrophied hepatic stellate cells for their transcriptomic characterization from CDAHFD mice liver

**DOI:** 10.1101/2025.05.19.654829

**Authors:** Marion Heckmann, Nour-El-Houda Djerir, Pierre-Henri Commere, Julien Fernandes, Guillaume Sarrabayrouse, Pascal Bigey, Virginie Escriou, Céline Hoffmann

## Abstract

Hepatic stellate cells (HSC) are known for their major role in hepatic fibrosis. Recently, it has been shown that different subpopulations of HSC co-exist during fibrogenesis and play different roles in the establishment of fibrosis. We previously highlighted, in murine model and human biopsies, a specific subpopulation of hypertrophied HSC (hypHSC) which exhibit exacerbated retinoid droplets and were closely associated to collagen fibers. The present study describes the purification protocol of hypHSC developed from a murine model of metabolic liver fibrosis. Liver dissociation followed by density gradient and fluorescence assisted cell sorting allowed us to obtain higlhly pure hypHSC preparations. Then, a transcriptomic analysis (bulk RNAseq) of hypHSCs versus quiescent HSCs purified from healthy mouse liver, was performed. This showed that hypHSCs molecular signature differs from HSC subtypes already described in the literature, with a “hybrid” profile of both inflammation and matrix remodeling. Our study highlights that a phenotype-to-molecular approach can provide complementary elements to single-cell molecular approaches.

## Introduction

Hepatic fibrosis, which corresponds to the excessive accumulation of extracellular matrix proteins in the liver parenchyma, is a major consequence of chronic liver diseases such as viral hepatitis, alcoholism and metabolic dysfunction-associated steatohepatitis (MASH, previously NASH)^1^. Hepatic fibrosis is a critical step in the course of the disease, as it can evolve into cirrhosis and hepatocellular carcinoma, which is one of the most serious cancers*^2^*. No specific treatment for hepatic fibrosis and its consequence is available; the only therapy is liver transplantation when the disease has reached its endpoints.

Among the liver cells, hepatic stellate cells (HSCs) are the major cells responsible for the synthesis of the extracellular matrix, mainly composed of collagen in liver tissue, during fibrosis^3^. Upon liver injury, HSC are known to become activated and transdifferentiate into proliferative, fibrogenic myofibroblasts and it is now well established that they are a central driver of fibrosis in experimental models and in human liver injury*^4^*. Recent studies based on single cell RNA sequencing (sc-RNAseq) highlighted the co-existence of different HSC subtypes during fibrogenesis with different activated ‘profiles’ such as inflammatory, proliferative or extracellular matrix producing, revealing the complexity of HSC activation and the phenotypic plasticity of HSC ^5-8^. These different subpopulations have been identified in different mouse models of fibrosis associated with NASH, CCl_4_ exposition or cholestasis (BDL model) as well as in human.

Moreover, the existence of two subpopulations of quiescent HSCs (qHSCs) expressing different genes depending on their location in the hepatic lobule, central vein-associated HSCs (CaHSCs) and portal vein-associated HSCs (PaHSCs), has recently been revealed in murine model of fibrosis as well as in human liver ^5, 9^.

In this context, we previously discovered in 4 murine models of liver fibrosis (diet- or chemically-induced) as well as in human liver biopsies with hepatic fibrosis due to different etiologies (alcoholism, NASH or hepatitis C) the existence of a specific phenotype of hepatic stellate cells associated to collagen accumulation zones^10^. These cells exhibit a hypertrophied phenotype, with exacerbated accumulation of cytosolic droplets containing retinoids as we demonstrated that they emit vitamin A-specific fluorescence. We called these cells hypertrophied hepatic stellate cells (hypHSC) and demonstrated that they express both markers of quiescent (desmin, cRBP1) and activated HSC (α-SMA)^10^. Using a combination of fluorescent and non-linear microscopies and the specific intrinsic retinoid droplet fluorescence, we were able to identify hypHSC in murine and human liver slices and determine their association with collagen deposition. In particular, we demonstrated a significant positive correlation between the stage of hepatic fibrosis and HSC hypertrophy in a cohort of obese patients with hepatic fibrosis.

We therefore hypothesize that hypHSC constitute one of the hepatic stellate cell subpopulations that could play a direct or indirect role in liver fibrosis. Further characterization of these cells, such as analysis of their gene expression, appears necessary. So far, hypHSC have been identified by microscopy analysis of murine liver slices and human liver biopsies with fibrosis. The next step is to produce a purified preparation of these cells.

The aim of this paper is to describe the experimental strategy developed to obtain a purified preparation of hypHSC allowing transcriptomic analysis. For this work, we have chosen the CDAHFD (choline deficient amino acid defined high fat diet) murine model we had previously used, since it is a murine model of MASH that recapitulates all the stages of the disease with steatosis, inflammation, progressive fibrosis associated with HSC hypertrophy with a very high reproducibility.

The isolation strategy consists of two main steps: 1) hepatic dissociation and enrichment in HSCs by density gradient followed by 2) fluorescence cell sorting (FACS) thanks to hypHSC or qHSC intrinsic fluorescence. We also describe a first characterization of hypHSCs at protein-level marker expression, followed by transcriptomic analysis (bulk RNA-seq), comparing their gene expression profile with that of qHSCs.

## Results

### 1. Enriched hypHSC preparation from CDAHFD mouse liver

#### 1.1. Protocol design

Our starting point for hypHSC isolation, relied on the well-described isolation protocol described by Mederacke *et al.* (figure 1). This method consists of *in situ* dissociation of the liver by enzyme perfusion, followed by excision of the liver to recover the cells as a suspension and by density gradient for separation of high-lipid-content cells ^11-13^ (Figure 1).

**Figure1:**
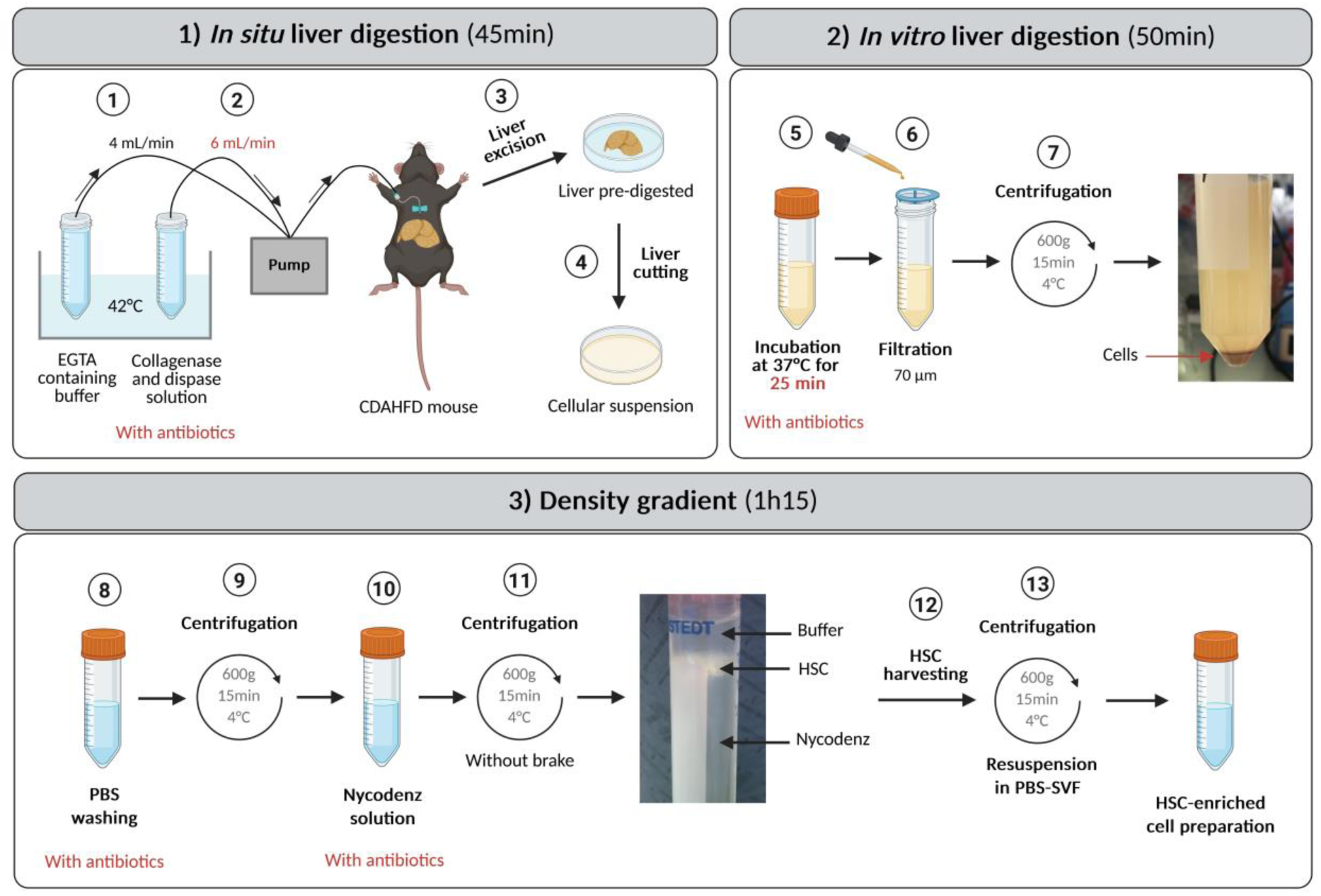
Schematic representation of the steps involved in the purification of hypertrophied HSCs from CDAHFD mice. Changes in conditions compared with the purification of quiescent HSCs from SD mice are highlighted in red. 1) Perfusion of the inferior vena cava by insertion of a 21G needle connected to a pump with perfusion buffer (containing EGTA) at 4 mL/min. 2) Perfusion of the liver with a buffer containing enzymes (collagenase and dispase) buffer and antibiotics (penicillin, streptomycin and gentamicin) and increased perfusion rate to 6 mL/min. 3) Excision of the digested liver and transfer to a Petri dish containing buffer with enzymes and antibiotics. For experiments requiring sterile conditions, manipulate within a biosafety cabinet from this stage onwards. 4) Cutting of the liver to release the cells. 5) Incubation of the resulting cell suspension at 37°C for 25 minutes to continue *in vitro* digestion. 6) Filtration of the cell suspension through a 70µm-filter to remove undigested liver pieces. 7) Centrifugation to pellet the cells and to remove enzyme buffer. 8) Rinsing with PBS containing antibiotics. 9) Centrifugation to pellet the cells and remove PBS. 10) Mixing of the cell suspension with a Nycodenz solution. 11) Centrifugation to form the density gradient to separate HSCs, thanks to their low density, from other liver cells. 12) Harvesting HSCs floating on the gradient. 13) Centrifugation to remove buffer and resuspend the cells in PBS-SVF 1%. Created in BioRender. Hoffmann, C. (2025) https://BioRender.com/im0zs9r

In CDAHFD-murine model, hypHSCs start to be detected from 6 weeks of CDAHF diet (see Figure S1). At this point, the liver exhibits stage F1 fibrosis and stage 3 steatosis, requiring adaptation of the protocol to this collagen- and lipid-laden pathological microenvironment. Isolation of hypHSCs was performed from the liver of CDAHFD-fed mouse for at least 9 weeks of diet, since hypHSC content being stable from this time point. Quiescent HSCs, that were considered as control cells, were isolated from the liver of mouse fed with a standard diet (SD), between 6 and 10 months old to recover sufficient HSCs, as qHSC level increases with aging.

Firstly, the conditions of CDAHFD-mouse liver digestion were adapted from the qHSC isolation protocol, by increasing *i)* the enzyme concentrations (x1.5 for collagenase, x2 for dispase), *ii)* the perfusion flow rate (from 4 to 6 ml/min), *iii)* the digestion buffer volume (x1.5) and *iv)* the post-perfusion incubation time (+ 5 min). Secondly, the centrifugation steps were adjusted by resuspending the cells in PBS for washing before resuspending them in GBSS/B buffer for the gradient step, as we observed a loss of cells when washing in GBSS/B directly. Remarkably, after the first centrifugation step, we observed for CDAHFD-mouse liver cell suspension the formation of a fat layer on the surface of the supernatant that is not observed for SD-mouse liver cell suspension. This is due to the steatotic state of CDAHFD-mouse liver. After this first centrifugation step, the layer of fat was carefully removed before continuing the experiment.

Another major modification to the protocol was the addition of antibiotics (Penicillin/Streptomycin/Gentamycin) in each buffer since systematic bacterial contamination was observed in cell preparations from CDAHFD-mice liver, whereas this was never the case from SD-mice liver, despite the same precautions for sterility being taken. We hypothesized that contamination of CDAHFD preparation was due to an endogenous contamination of CDAHFD liver. Some authors described that under fat diet, an intestinal bacterial translocation may occur ^14, 15^. To ensure this hypothesis, we assessed the bacterial load in the liver and feces of CDAHFD- and SD-mouse using classical and real-time PCR amplification of the 16S rRNA gene. We observed a decrease in bacterial load in CDAHFD-mice feces and an increase in bacterial load in CDAHFD-mouse liver, as compared with samples from control mice. These observations could be linked to a relocation of bacteria from the gut to the liver, suggesting a phenomenon of bacterial translocation in our model (See Figure S2).

#### 1.2. Cell characterization after the gradient step

After density gradient, 6.4 ± 4.1 x 10^6^ cells (n=53) were recovered from CDAHFD-mice liver (9-15 weeks of diet) vs. 2.9 ± 2.0 x 10^6^ cells (n=16) from SD-mice liver (5-10 months old). Thus, on average, 2.2 times more cells are recovered from the liver of CDAHFD-mice than from SD-mice.

One of the specific characteristics of HSCs is that they contain cytoplasmic droplets rich in retinoids. These retinoids emit a specific endogenous fluorescence in the near UV-violet range, with excitation between 365 and 405 nm and fluorescence emission from 380 to 500nm ^16, 17^. Previously, we highlighted on CDAHFD-mouse liver sections, that hypHSCs exhibit specific and broad fluorescence properties with a fluorescence excitation in the UV-visible spectrum and emission throughout the visible spectrum (400-600nm). Notably, hypHSCs retain the fluorescence properties in the UV-violet range like qHSC (λ_exc_=365nm and λ_em, max_=390 nm) and after excitation at 488 nm, emit an additional intense signal in the green range (500-550nm), while this is not the case for qHSCs, which are not excitable beyond 405 nm ^10^.

In this study, we used these specific fluorescence properties to identify hypHSC and qHSC in cell preparations. We qualified the natural fluorescence of qHSC (λ_exc_=365nm, λ_em, max_=390 nm) as “blue” fluorescence and the specific hypHSC fluorescence in the green range (λ_exc_=488nm, λ_em, max_ =525nm) as “green” fluorescence. Therefore, qHSC are blue+/green- and hypHSC are blue+/green+ (Figure 2.A). As shown in Figure 2.B, after the gradient, the cell preparation from CDAHFD-mouse liver, contained approximately 1/3 hypHSC (blue+/green+) as well as other cells containing retinoids (blue+/green-) that may be remaining qHSC and also a majority of non-fluorescent cells. The preparation from an SD-mouse liver did not contain hypHSCs, only a large majority of qHSCs (blue+/green-) and a few non-fluorescent cells.

**Figure 2.**
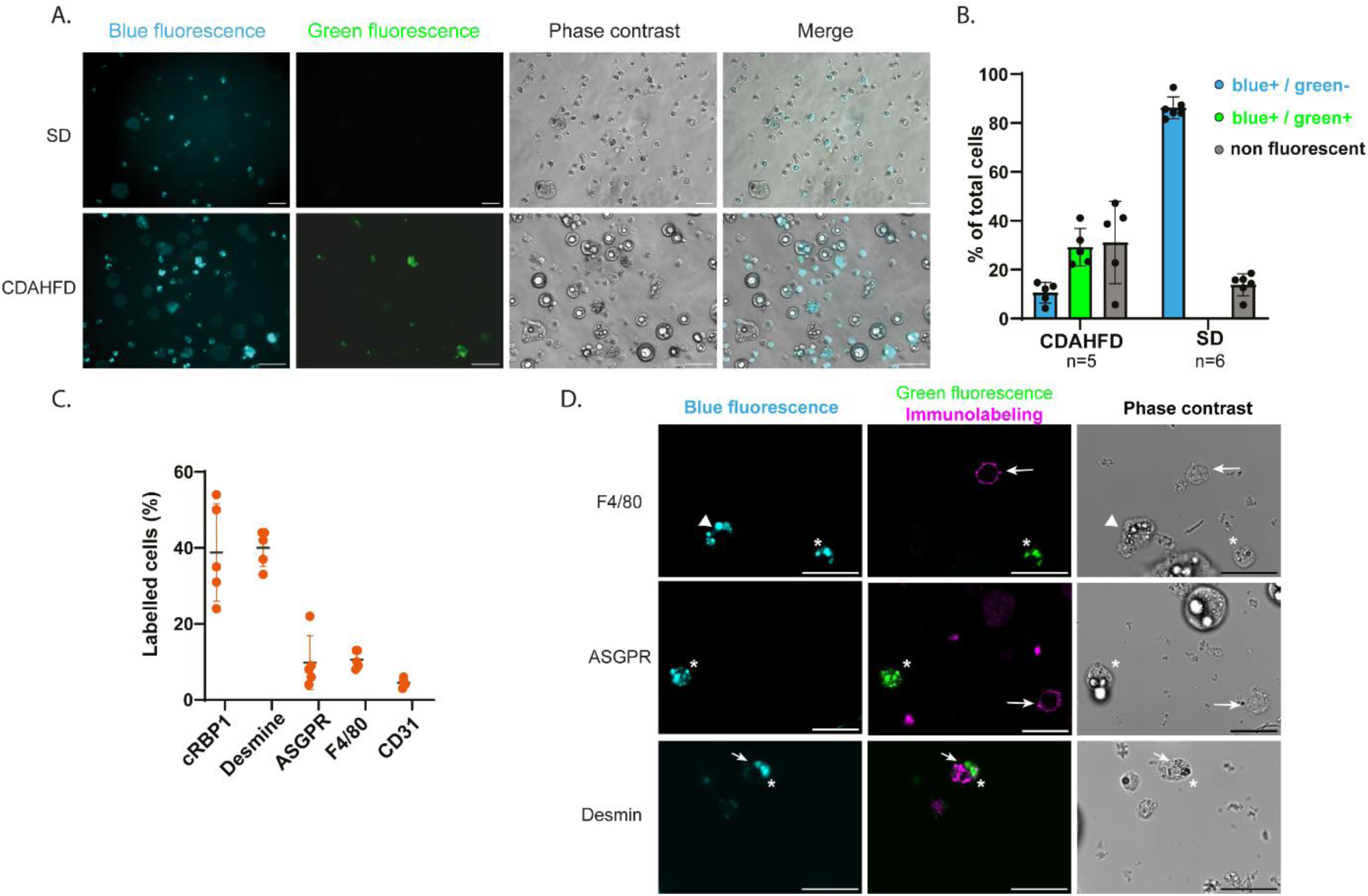
Characterization of cells obtained after density gradient. A) Representative microscopy images of cell preparation from CDAHFD- and SD-liver. Acquisitions under UV-Vis excitation for blue fluorescence (λexc=365nm λem >390nm), under excitation at 488 nm for green fluorescence (λexc=488nm, λem =500-550nm), phase contrast illumination, and merge of images. Scale bar: 50 µm. B) Percentage of fluorescent cells from CDAHFD- and SD-mouse liver counted from microscopic images, n(CDAHFD)=5 and n(SD)=6 independent experiments. C) Repartition of hepatic cell population in CDAHFD cell preparation using cytometry analysis. Percentages of labelled cells were obtained after immunolabeling of cells with typical markers of HSC (cRBP1 and desmin, n=5), hepatocytes (ASGPR, n=5), macrophages (F4/80, n=5) and endothelial cells (CD31, n=4). D) Representative microscopy images of cell preparation after density gradient from CDAHFD-liver after immunostaining with ASGPR, F4/80 or desmin antibodies (magenta, λexc= 638nm, λem= 650-690 nm) and acquisitions for blue fluorescence (λexc= 405 nm, λem= 425-465 nm, cyan) and green fluorescence (λexc= 488 nm, λem= 505-535 nm, green). Arrowhead: qHSC, Star: hypHSC, arrows: cells with positive corresponding immunolabelling. Scale bar=25 µm.

The composition of cell population from CDAHFD-mouse liver was further analyzed by flow cytometry after immunolabeling with antibodies specific to liver cells (CD31, endothelial cells; ASGPR, hepatocytes; F4/80, Kupffer cells; desmin and cRBP1, HSCs). As shown in Figure 2.C, we found 40 ± 4,8% of cells that were desmin+, and 38,8 ± 12,7% that were cRBP1 +, while the other antibodies did not label more than 10% of cells, confirming the enrichment of the preparation in HSC. Fluorescence microscopy imaging allowed us to better characterize the cell population obtained after density gradient (Figure 2.D). qHSC were identifiable thanks to blue fluorescence contained in cytoplasmic droplets and hypHSC exhibited, as previously described, blue and green fluorescence in cytoplasmic droplets. Immunolabeling of ASGPR and F4/80 was observed around cell membrane of non-fluorescent cells and never colocalized with cells containing fluorescent droplets. Unlike ASGPR or F4/80, desmin immunolabeling was colocalized with cells containing blue and blue/green cytoplasmic droplets corresponding to qHSC and hypHSC, respectively (stars and arrowheads on figure 2.D).

Thus, as hypHSC represented only 30% of the cell population obtained after gradient from CDAHFD-mouse liver, which was not sufficient for further characterization of these cells, an additional purification step was essential. To obtain a highly pure hypHSC preparation, we used fluorescence-assisted cell sorting (FACS), taking advantage of the specific fluorescence properties of these cells, thus allowing sorting to be carried out without requiring any exogenous labeling.

### 2. Sorting of hypHSC

#### 2.1. Sorting strategy

The cell preparation obtained after density gradient contains a mixture of cells of varying size and granularity, and probably cell debris. Cytometric analysis for both parameters (FSC and SSC) was not sufficient to identify with certainty the position on the cloud corresponding to the cells. Cell labelling with SYTO16 allowed us to determine the position of intact cells cloud on the FSC-SSC dot-plot (See Figure S3). Thus, after an initial selection of intact cells on the FSC-SSC dot-plot, the cells were analyzed according to SSC and green fluorescence (λ_exc_=488 nm, λ_em_=513/26 nm) or blue fluorescence (λ_exc_ = 405 nm and λ_em_ = 425-475 nm) (Figure 3.A). We then observed, for both CDAHFD and SD cell preparations, the presence of a blue-fluorescing cell population corresponding to qHSC. And as expected, a population of cells emitting green fluorescence was found in the CDAHFD cell preparation but not in the SD cell preparation, so we assumed this part of the cloud to be hypHSC (fig 3.A).

**Figure 3.**
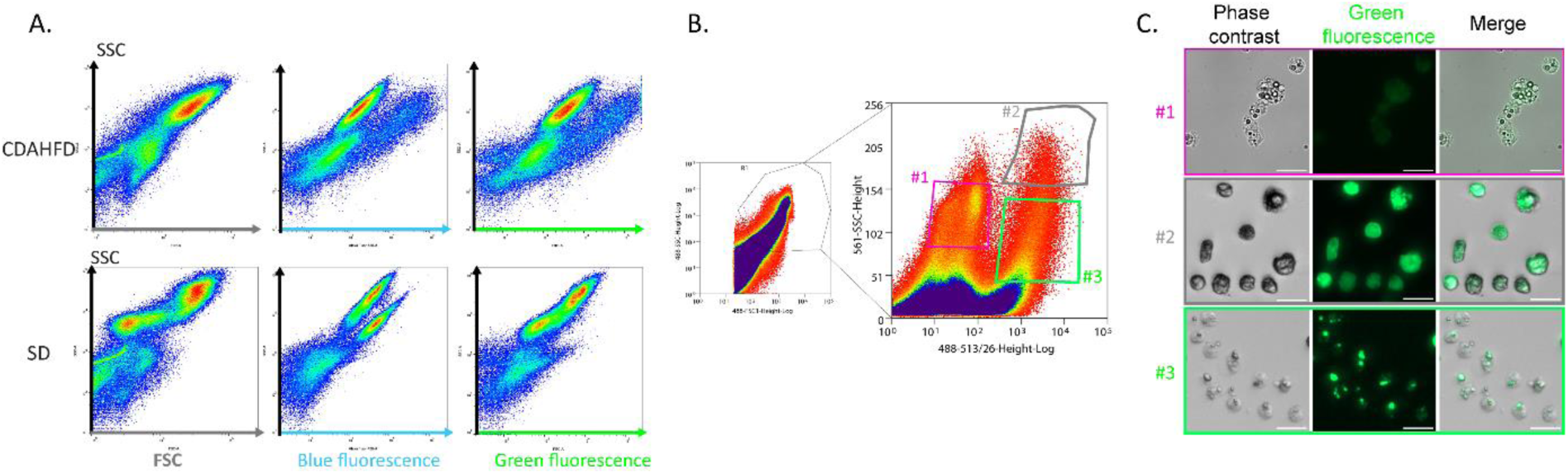
Sorting strategy for hypertrophied HSC using fluorescence-activated cell sorting. A) Cytometric analysis of cell preparations obtained from CDAHFD- and SD-mice liver after density gradient. Dot plot of the CDAHFD and SD cell preparation analyzed for FSC/SSC (left panels), SSC and blue fluorescence (λexc= 405 nm and λem,max= 450 nm; middle panels), SSC and green fluorescence (λexc= 488 nm and λem,max= 530 nm; right panels). B) FACS gating strategy for hypHSC purification: SSC-FSC dot plot allowing to determine the first gate for cells, and dot plot for SSC (linear)–green fluorescence to determine the gate for hypHSC sorting. #1, #2, #3 correspond to 3 gates selected to characterize the ideal gating strategy for hypHSC purification. C) Phase contrast and green fluorescence (λexc=488nm, λem =500-550nm) microscopy images corresponding to the gates #1-#3 on the SSC-green fluorescence dot plot. Bars= 25µm.

Starting from these analyses, the hypHSC sorting strategy was based on the unique green fluorescence of hypHSC preceded by a succession of sorting gates. Sorting was carried out on an Astrios equipped with a 100 µm nozzle, allowing the sorting of cells from liver cell preparation (containing large cells) with a relatively low pressure flow regarding the fragility of hypHSC. First, cell selection was performed on the FSC–SSC dot-plot to eliminate cellular debris; then, a selection of single cells was made to eliminate doublets; finally, the cell cloud was selected on green fluorescence signal to recover hypHSCs (Figure 3.B). To select hypertrophied HSCs more accurately, the SSC parameter of the green fluorescence cloud of HSC had to be expressed on a linear scale, to obtain a more spread-out distribution of points. The area corresponding to the fluorescent cells was then selected more precisely (figure 3.B, and see Figure S4).

Different gates for recovering the cells were determined on the SSC-lin/green-fluorescence plot and the cells morphology was analyzed by fluorescence microscopy to determine the best gate for hypHSC sorting (Figure 3.C). As expected, cells from gate #1 characterized by low fluorescence signal (between 10^1^ and 10^2^) exhibited a low fluorescence signal detectable by fluorescence microscopy (figure 3.C) while the cells from gates #2 and #3 are characterized by a high green fluorescent signal (between 2.10² and 10^4^), corresponding to hypHSCs. Gate #2 corresponded to highly fluorescent and granular cells. Microcopy observation of these cells revealed that their fluorescence signal was diffuse throughout their cytoplasm and phase contrast illumination showed poorly defined or unstructured droplets making the cells appear “opaque”. The cells from gate #3 presented a high green fluorescence and a lower granularity than cells from gate #2. Microscopy observations showed that the majority of cells recovered from gate #3 exhibited a green fluorescence signal arising from intact and well-defined droplets, observable under both fluorescence or phase contrast illumination (figure 3.C). We chose to harvest hypHSC with intact retinoid droplets, as this is the form in which we have previously identified them in CDAHFD-mice liver sections and human biopsies^10^. We then quantified the proportion of intact hypHSC (*i.e.* hypHSCs with intact retinoid droplets) and opaque hypHSCs (i.e. hypHSC with poorly defined retinoid droplets) in both gate#2 and gate#3. We established that gate#3 contained a majority of intact hypHSC with 56 ± 23 % (n = 4) against 41 ± 23 %, (n = 4) of opaque cells. In contrast, the proportion of cells from gate#2 was distributed such that 70 ± 12 % (n = 4) were opaque cells and 24 ± 10 % (n = 4) were intact hypHSCs. Therefore, for further experiments, the final gating strategy chosen for hypHSC sorting was the one corresponding to gate #3 (figure 3.B).

In parallel, qHSC were sorted, from a cell preparation derived from SD-liver, using the same cell sorter and following a succession of sorting gates similar to those of hypHSC, but with a final selection performed on the basis of SSC-log/blue fluorescence (λ_exc_=405nm; λ_em, max_= 455nm) as previously reported in the literature ^11, 12^.

#### 2.2. Cell characterization after the sorting step

At the end of the sorting step, 2.4 ± 1.1 x 10^5^ (n=30) hypHSCs were recovered from CDAHFD-mouse liver (9-15 weeks of diet) *vs* 4.4 ± 3.0 x 10^5^ (n=13) cells from a SD-mouse liver (5 to 10-month-old) (see Table 1). No significant difference was observed in the number of cells recovered depending on the diet time (9-19 weeks – see Figure S5). The sorting step recovered approximately 3.5 ± 1.1 % of hypHSC from the total CDAHFD-cell preparation obtained after density gradient.

**Table 1.** Number of cells obtained at the different stages of HSC purification. Cells were counted manually using Mallassez cells. Dead cells were quantified using Trypan Blue coloration. For CDAHFD-mouse liver, n=53 liver dissociations followed by density gradient were performed. Among them, 30 have been followed by FACS. Percentage of dead cells and hypHSC purity were quantified for 25 and 19 independent hypHSC sorted preparations respectively. For SD-mouse liver, n=16 liver dissociations followed by density gradient were performed. Among them, 13 have been followed by FACS. Percentage of dead cells and qHSC purity was quantified for 10 and 8 independent qHSC sorted preparations respectively.

| Diet | Average number of cells |  | % of cells sorted vs cells after gradient | % of dead cells | % of hypHSC or qHSC purity |
| --- | --- | --- | --- | --- | --- |
|  | after gradient | after sorting |  |  |  |
| CDAHFD (9-15 wk) | $6.4 \pm 4.1 \times 10^6$ (n=53) | $2.4 \pm 1.1 \times 10^5$ (n=30) | $3.5 \pm 1.1 \%$ (n=30) | $22 \pm 14\%$ (n=25) | $94.8 \pm 4.5 \%$ (n= 19) |
| SD (5-10 months) | $2.9 \pm 2.0 \times 10^6$ (n=16) | $4.4 \pm 3.0 \times 10^5$ (n=13) | $21.64 \pm 9.1 \%$ (n=13) | $12.2 \pm 8.7 \%$ (n=10) | $94.0 \pm 3.2 \%$ (n= 8) |

The hypHSC sorted preparations were very pure since 94.8 ± 4.5 % of the cells were hypHSCs with 22 % of mortality (Table 1). Considering that after density gradient the contamination by non-HSC cells was greater than 50%, cell sorting allowed the elimination of undesirable cells and represents a fundamental step for the further study of hypHSCs (Figure 4.C). In comparison, equivalent results in terms of purity and mortality were obtained for the preparations of qHSCs from SD-mouse liver, underlining that the sorting step is also important for qHSC study (figure 4.D). Importantly, the total duration of purification (dissociation followed by density gradient) and sorting had an effect on the viability of the hypHSCs obtained, as a 2-day purification increased significantly the percentage of hypHSC mortality (See Figure S5). Following cell sorting, we showed that green and blue fluorescence were indeed associated with well-defined droplets in hypHSCs, just as blue fluorescence was associated with retinoid droplets in qHSCs (figure 4.A-B).

**Figure 4.**
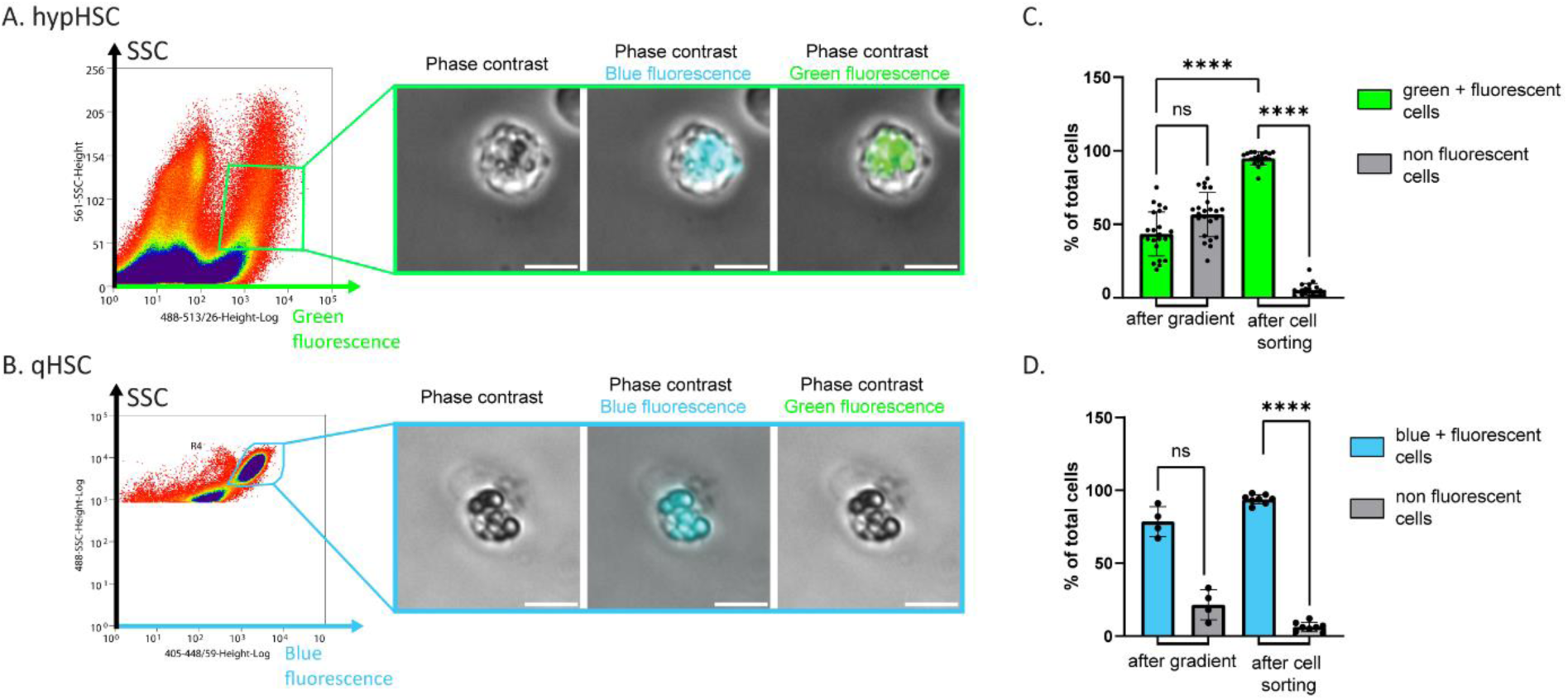
Morphology and characterization of sorted hypHSC and qHSC cell preparations. Representative dot plots with the gates used for sorting (A) hypHSC (top panel - SSC/green fluorescence) and (B) qHSC (bottom panel - SSC/blue fluorescence) and corresponding representative microscopy images of sorted hypHSC and qHSC. Images of microscopy performed under phase contrast illumination, and merge with green (λexc=488nm, λem =500-550nm) or blue fluorescence (λexc=365nm λem >390nm). Scale bar: 10 µm. C) Percentage of green fluorescent cells (hypHSC) and non-fluorescent cells in preparations obtained after density gradient and after sorting from 9 to 15 weeks CDAHFD-mice liver. Means ± standard deviation are shown, n(after gradient)= 23, n(after sorting)= 19, Kruskall-Wallis test. D) Percentage of blue fluorescent cells (qHSC) and non-fluorescent cells in preparations obtained after density gradient and after sorting from SD-mice liver. Means ± standard deviation are shown, n(after gradient)= 4, n(after sorting)= 8, One-way ANOVA test. *ns* p > 0.05; *p ≤ 0.05; **p ≤ 0.01, *** p ≤ 0.001, **** p ≤ 0.0001

#### 2.3. Stellate cell protein markers in sorted hypHSCs

Thanks to cell sorting, highly pure hypHSC and qHSC cell preparations were obtained and allowed us to examine the expression of stellate cell markers at the protein level by immunolabeling. Immunofluorescence study of sorted hypHSCs showed that they express desmin, cRBP1 and αSMA (figure 5.A) confirming what we previously showed by immunohistochemistry on CDAHFD-liver slices^10^. According to qPCR results, the level of αSMA expression in hypHSCs was equivalent to that of qHSCs (figure 5.B). Surprisingly, LRAT immunolabelling was low to not detectable, suggesting that hypHSCs have low LRAT expression that was confirmed by qPCR analysis (figure 5.B).

**Figure 5.**
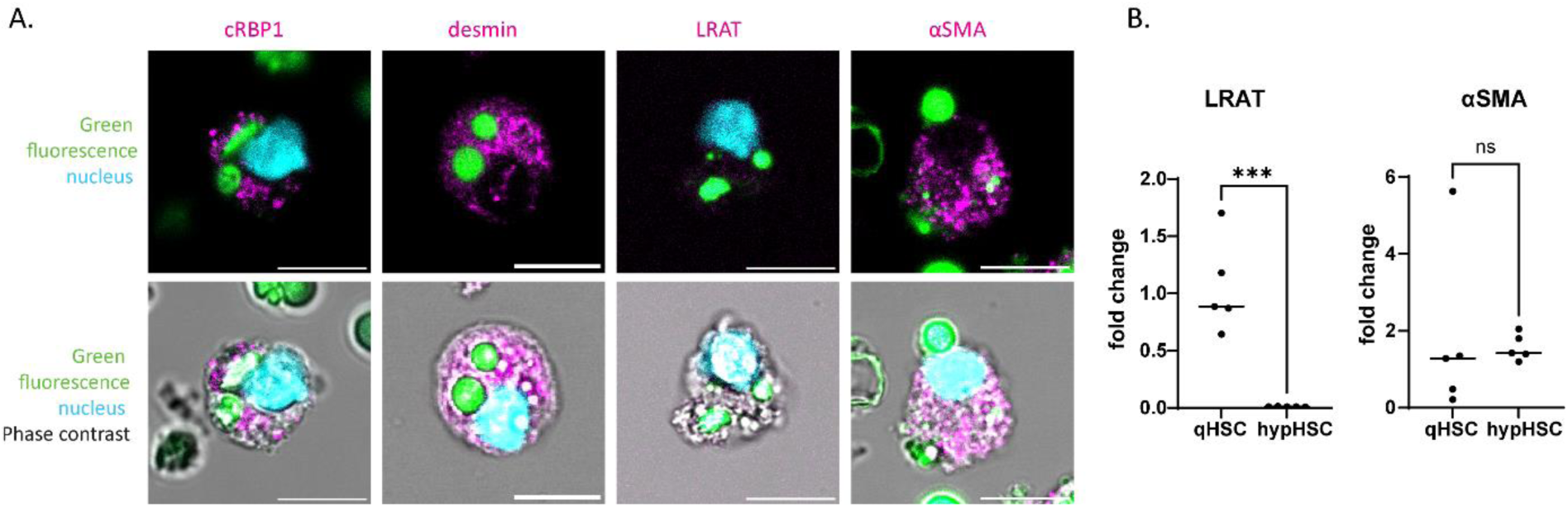
Analysis of stellate cell marker expression in hypHSC sorted cells. A) Representative images of cRBP1, desmin, LRAT and αSMA immunostaining (λexc= 638nm, λem= 650-690 nm, magenta) merged with “green” retinoid fluorescence (λexc= 488 nm, λem= 505-535 nm, green), nucleus (DAPI, λexc= 405 nm, λem= 425-465 nm, cyan) (top panels) and phase contrast illumination signal (bottom panels) acquired by confocal microscopy. Scale bar 10 µm. B) Relative expression of LRAT and αSMA in hypHSC vs qHSC analyzed by qPCR (statistical tests: for LRAT n=5, Unpaired t-test, for αSMA: n=5 Mann-whitney test, ns p > 0.05; *p ≤ 0.05; **p ≤ 0.01, *** p ≤ 0.001, **** p ≤ 0.0001).

Thus, we succeeded in purifying the hypHSC we previously identified by fluorescence microscopy on CDAHFD-mouse liver slice and human fibrotic liver biopsies from CDAHFD-mouse liver.

### 3. Bulk RNAseq analysis of hypHSC vs qHSC

As sorted hypHSC and qHSCs were obtained with 95% purity, this allowed us to further characterize hypHSCs from a molecular point of view. Although no difference was observed at the cellular level depending on CDAHFD time of diet, we decided to analyze sorted hypHSC from CDAHFD-mouse liver at restricted duration of 12 to 13 weeks of diet to limit possible molecular variability. Then, NGS sequencing of sorted hypHSC cells was performed with sorted qHSC as reference, bulk analysis allowed us to perform an in-depth analysis of gene expression.

#### 3.1. RNA extraction from sorted hypHSC and qHSC

The first important criterion before performing RNA sequencing is based on the quality of the extracted RNA, determined by RIN (RNA integrity number), used to produce the sequence library. As the RIN of sorted hypHSCs exhibited significant variability between samples, with RIN values ranging from 3 to 6 (See Figure S5), we therefore optimized the protocol by *i)* carrying out the different steps of cell purification (dissociation / gradient / sorting) on the same day, allowing lower cell mortality (See Figure S5), ii) recovering sorted cells directly in lysis buffer (used for RNA extraction) with an optimal number of cells, iii) using a RNA extraction kit adapted to low cell number. These optimizations increased the RIN of RNA extracted from sorted hypHSCs with a maximum value of 7.3. Then, for RNAseq analysis of hypHSCs, we selected, among 36 samples obtained from the liver of 19 CDAHFD mice (12-13 weeks of diet), 5 preparations with the highest RINs (mean 6.45). In parallel, 5 preparations of RNA extracted from sorted qHSCs were also selected (mean RIN 7.14 with no difference in mean RIN of selected hypHSC samples).

#### 3.2. Bulk RNAseq analysis of sorted hypHSC versus sorted qHSC

Once samples with satisfying RNA quality were selected, non-strand oriented libraries were prepared as described in the material and method part. As the RIN of the selected hypHSC samples was slightly lower than that of the qHSCs, we carefully analyzed the quality of generated librairies. Analyses of sequencing data quality, reads repartition (e.g., for potential ribosomal contamination), inner distance size estimation, genebody coverage, strand-specificity of library were performed using FastQC v0.11.2, Picard-Tools v1.119, Samtools v1.0, and RSeQC v2.3.9. These first quality controls were assessed using MultiQC including mapping statistics from STAR and read assignments from FeatureCounts.

A total of 16,066 genes were identified on the 10 samples (5 hypHSC, 5 qHSC) during sequencing. Among the genes identified, some genes described as HSC marker genes in the literature ^18, 19^ were found and expressed by hypHSCs and qHSC in terms of FPKM (Fragments Per Kilobase per Million mapped fragments) such as vimentin, desmin, acta2, and several collagens (col1a1, col1a2 and col3a1) confirming the stellate cell status of hypHSCs (See Figure S6).

Analysis of differentially expressed genes (DEGs) revealed 6,926 DEGs between the hypHSC and qHSC populations, with 3,369 genes overexpressed by hypHSCs compared with qHSCs and 3,557 genes underexpressed by hypHSCs compared with qHSCs (fig 6.A). This result points out a marked difference in gene expression between the two HSC populations.

**Figure 6.**
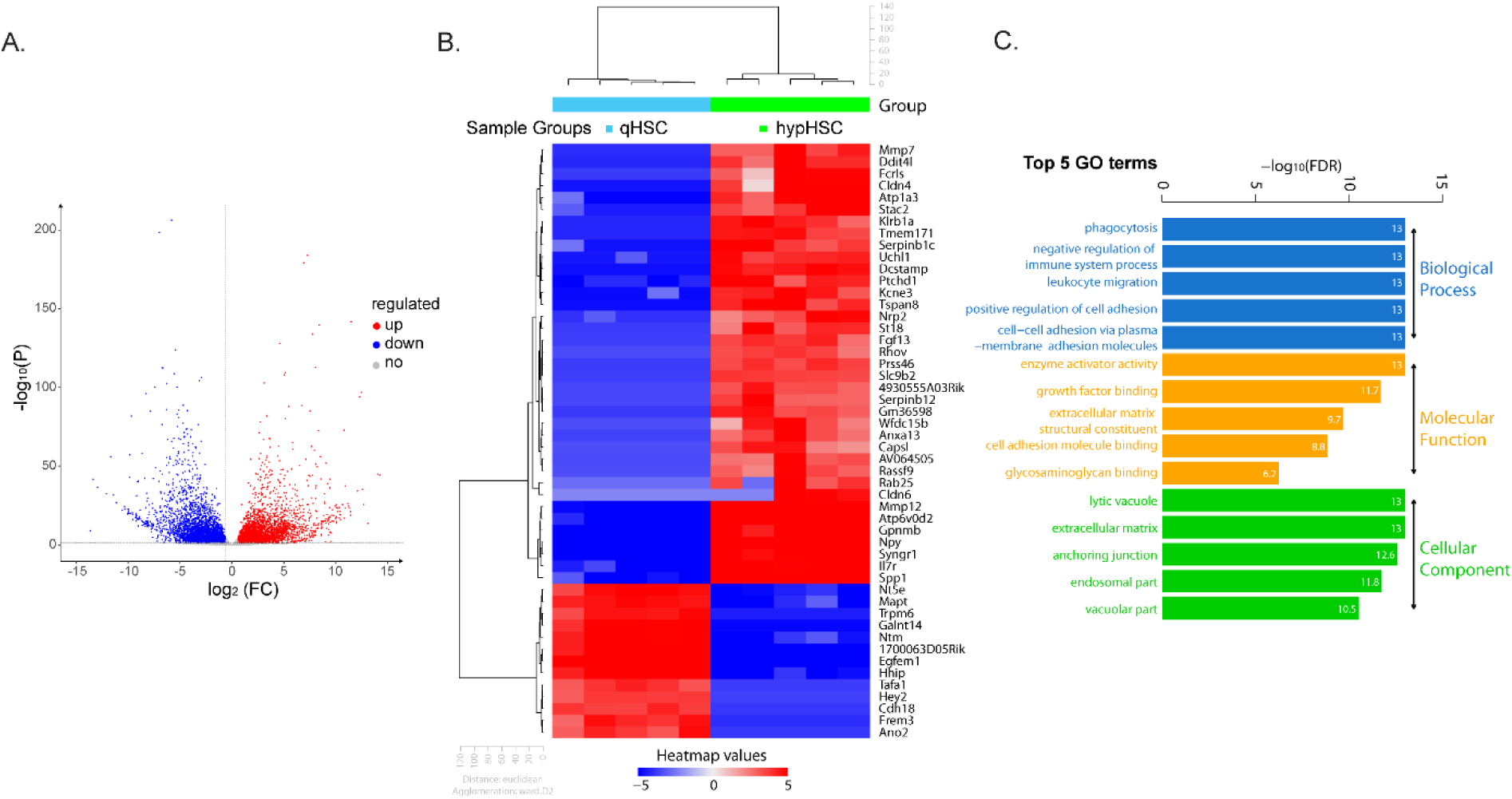
RNAseq analysis of sorted hypHSC and qHSC. A) Volcano plot depicting differentially expressed genes (DEGs) between hypHSCs and qHSCs. Blue: downregulated; grey: unregulated; red: upregulated. B) Heatmap and unsupervised hierarchical clustering of the top 50 DEGs, C) Top 5 gene ontology (GO) analysis terms for biological process, molecular function and cellular component.

By identifying the 50 most DEGs, we defined the molecular signature of hypHSCs (fig 6.B). Among the panel of HSC-specific genes, we observed a strong modification of their expression. Thus, some quiescent HSC-specific genes appeared downregulated (such as sparc, dcn, csrp2, bambi…) and others upregulated such as adipor1, plin2, vimentin and pparg (See Table S1). Interestingly, the “gold standard” markers of quiescent HSCs such as lrat (log 2 FC 6.05), gfap (log2 FC 4.37) and desmin (log2FC 5.44) appeared to be down regulated, although the latter was detectable at protein level on sorted hypHSCs (fig 5.A). Moreover, most of the genes known as markers of activated HSC did not appear deregulated, such as acta2, collagens (col1a1, col1a2, col3a1), notch 3 and timp1 ^20^ (See Figure S6 and Table S1).

To better characterize the expression differences between hypHSCs and qHSCs, a gene ontology (GO) analysis was performed to identified DEGs into 3 categories: biological process, molecular function and cellular component (fig 6.C). This analysis suggested an interaction of hypHSCs with the immune system, with cell adhesion functions (down-regulated), with phagocytosis/endocytosis (upregulated), and an involvement in the regulation of ECM composition. Concerning the regulation of extracellular matrix composition, MMP12 and MMP7 appeared to be upregulated and within the first 50 DEGs, while laminin and fibronectin appeared to be down-regulated, suggesting a role of hypHSCs in modifying the balance of extracellular matrix synthesis and production.

Moreover, some genes involved in inflammation and regulation of immune system process, such as IL7r (in the TOP 50 DEGs) and TNF-α, appeared to be significantly upregulated.

Finally, genes associated to phagocytosis/endocytosis process were also upregulated, such as Trem2, CD36 (phagocytosis), CD68 and CD63 (endocytosis) or cathepsins.

Taken together, these results suggest that hypHSCs cannot be defined as “classical” activated HSCs or “quiescent” HSCs. They may suggest that hypHSCs exhibit a hybrid profile involving multiple pathways associated with the fibrogenesis process that does not correspond to the HSC subtypes already described in the context of liver fibrosis.

## Discussion

In this work, we have described the strategy for isolating hypHSCs, a specific population of HSCs associated with hepatic fibrosis. Although the protocol for HSC isolation is widely described for healthy or fibrotic murine liver (mainly CCl_4_ model) ^21, 22^, there is only a few papers on HSCs isolation from fatty and fibrotic liver. To our knowledge, only one study has performed HSCs isolation from a CDAHFD liver, after 3 weeks of diet. In these conditions, the liver does not yet present fibrosis but already steatosis^23^. In the same way, one paper reports the isolation of HSCs from mice fed a "Western diet" for 12 weeks^7^ without specifying the influence of steatosis on HSC purification or HSC purity. Thus, the hypHSC isolation protocol we describe here, based on the CDAHFD model and characterized in a large number of mice (n = 19), can be applied to studies on HSC in a context of metabolic hepatic fibrosis. Our study also highlights the importance of cell sorting after the density gradient, as FACS yielded an ultra-pure hypHSC fraction (95%) from an enriched hypHSC fraction (43%) after the density gradient.

Our results report an augmentation of bacterial load in the liver of CDAHFD-mice compared to healthy mice. These observations are consistent with literature data describing a link between MALFD/MASH and disruption of the intestinal microbiota. Indeed, a change in the gut microbiota following a high-fat diet could be associated with an increase in the permeability of the intestinal barrier, thus allowing the translocation of intestinal bacteria towards the liver *via* the portal circulation. For example, mice fed a high-fat (HF) or choline- and methionine-deficient (MCD) diet have an abnormal composition of the intestinal microbiota, leading to a decrease in the thickness of the mucous layer of the intestinal barrier and a redistribution of tight junctions forming proteins^15, 24^. Increased intestinal permeability and impaired tight junctions have also been observed in patients with NAFLD ^25^. The onset of intestinal bacteria in the liver causes hepatic inflammation as bacteria and their metabolites constitute PAMPs (Pathogen-Associated Molecular Pattern) that can bind to TLR-4 and TLR-9 receptors on Kupffer cells^26,27^. The resulting inflammation can lead to the formation of steatohepatitis in the context of NAFLD. However, the cause and effect relationship between disturbance of the gut microbiota and MAFLD/MASH is unclear and two hypotheses are considered: disturbance of the microbiota can promote the formation of NASH by causing liver inflammation, or it is hepatic steatosis that causes a disruption of the gut microbiota *via* changes in liver secretions carried by bile. Our data showed that the purification of HSCs from mice subjected to a CDAHF diet, requires precautions such as adding antibiotics to the different solutions used. In addition, these results indicated that the use of CDAHFD model could be interesting to study microbiota and bacterial translocation in the context of metabolic liver disease.

In this paper, we showed that the characteristic fluorescence properties of hypHSCs in the green range allowed their specific detection during isolation from an enriched fraction of HSCs obtained after density gradient. Using the fluorescence properties of retinoid droplets in the blue range is a well-known and widely used strategy for the purification of quiescent HSCs ^11-13^ but to our knowledge, our study is the first that uses distinct intrinsic fluorescent properties to isolate a particular HSC population in a fibrotic context. The different fluorescence properties between hypHSC and qHSC retinoid droplets may be due to a different composition in term of triglycerides or fatty acids or even a difference in lipid bilayer morphology, as it has been demonstrated to occur during HSC activation ^28-30^. Moreover, an increase of free cholesterol content in HSC has been shown in murine models of nonalcoholic steatohepatitis^31^. Then, further study on the content of hypHSC lipid droplet could be of interest to understand the dynamic of retinoid droplets that occurs upon HSC hypertrophy in the context of hepatic fibrosis and for such a study, isolated hypHSC are necessary.

While these specific properties allowed us to distinguish hypHSCs from other cells, they seemed to add difficulty for their characterization based on fluorescence analysis methods. Therefore, all steps of the process were monitored by cell microscopy observation to ensure that they contain intact and fluorescent droplets. However, this fluorescence broad spectrum may induce a high “noise” signal that has to be considered and could require spectral acquisition and compensation analysis for further study (See Figure S7).

Bulk RNAseq analysis of hypHSCs allowed us to perform in-depth analysis of gene expression and to determine a difference of gene expression for 6,926 genes between hypHSCs and qHSCs. We showed that hypHSCs resemble neither qHSCs nor activated HSCs, as typical aHSC genes such as collagens or sparc and crsp2 did not appeared deregulated ^18^. It is now well known that the paradigm of “quiescent” or “activated” cellular states for HSCs no longer fully reflects the heterogeneity of HSC subtypes, as scRNAseq studies shed light on the dynamics of HSC-myofibroblast transition in response to liver injury. However, none of these studies identified hypHSCs, possibly due to the strategy chosen to isolate HSCs, based on transgenic mice expressing a reporter gene under a HSC-specific promoter such as that of LRAT or PDGFR-β ^5, 21^. Such a strategy, rather elegant for the isolation of cells, is likely to induce a bias in the study since it assumes that HSCs are characterized by a unique marker, the gene associated with the promoter, contrary to the strategy we used here, based on the specific and endogen fluorescence of hypHSCs in the green range.

However, some scRNAseq studies based on a non-parenchymal hepatic fraction (*i.e.* after gradient) or on sorted HSCs based on ‘blue’ fluorescence properties in several fibrotic contexts (CCl_4_, NASH or cholestatic murine model) should have recovered hypHSCs simultaneously with other HSCs subtypes. For example, we found that hypHSCs could correspond to the cluster of activated HSCs highly expressing genes associated with leukocyte activation and immune regulation such as Slpi, Cd74 and Dmkn described in Krenkel’study ^6^. By comparing the genes expressed by hypHSCs we hypothesized that they could correspond to cluster #7 of Rosenthal’study which is part of the 3 HSC clusters not described in details^7^.

Indeed, our transcriptomic data showed that hypHSCs underexpressed certain typical HSC genes such as LRAT, PDGFR or desmin, compared to qHSCs. Our data suggest that they have a hybrid profile with expression markers specific of phagocytosis and inflammation, while expressing some markers of extracellular matrix remodeling.

Then, we demonstrate here that by adopting an unusual “top-down” approach (from the specific phenotypic characteristics of hypHSCs down to their gene expression), we were able to purify these cells and characterize their molecular signature that differs from already described HSC subtypes in the fibrosis context.

Based on this study, further investigations may help elucidate hypHSCs status among the different HSC subpopulation that coexist during hepatic fibrosis and better understand their role in the mechanisms of fibrosis.

## Material and methods

### 1- Animals and experimental design

C57BL/6 mice (Janvier Labs, France) were housed in cages (5 per cage) and were kept at 23 +/-3 °C with a 12:12 h light/dark cycle. Male 6-month-old (former breeders) or 6-week-old mice were fed a standard diet (SD, Specific Diet Services, England 841201, vitamin A content 8000 IU/kg) for up to 6 months. After 5 days of acclimation, male 6-week-old mice were fed a choline-deficient, 0.1% methionine, high-fat diet (CDAHFD, Safe Diets, France U8958P Version 0247, vitamin A content 5280 IU/kg) for up to 20 weeks. Mouse weight, behavior and appearance were monitored weekly. All animal experiments have been approved by the Ethics Committee for Animal Experimentation and authorized by the French Ministry of Higher Education and Research (APAFiS #33532-2022030115518551 v3 and #46428-2023092921282299 v4).

### 2- HSC isolation

#### *In situ* enzymatic dissociation of the liver

Mice (SD or CDAHFD fed) were anesthetized with a mixture of ketamine/xylazine (180/20 mg/kg) by intraperitoneal injection. After checking for deep anesthesia by absence of paw withdrawal, a laparotomy is performed followed by lifting of costal plastron quickly after respiratory arrest. Perfusion of the liver through the inferior vena cava was then performed with a 21G needle (Venofix® A, B. Braun) connected to pre-heated (42°C) perfusion buffer (25 mM HEPES, 1.2 mM KH_2_PO_4_, 2.3 mM KCl, 86 mM NaCl, 0.5 mM EGTA, pH 7.6) at a rate of 4mL/min. Then the portal vein was cut and perfusion continued with 45 ml of perfusion buffer at a flow rate of 4 ml/min. Liver perfusion was followed with 25 mL (SD mice) or 50 mL (CDAHFD mice) of pre-heated (42°C) digestion buffer (25mM HEPES, 2.3 mM KCl, 86 mM NaCl, 5mM CaCl_2_, pH 7.6) containing Collagenase type I (Gibco 17100-017) at 205 U/mL for (SD mice) or 375 U/mL (CDAHFD mice) and Dispase (Gibco 17105-041) at 1U/mL (SD mice) or 2 U/mL (CDAHFD mice) at a rate of 4mL/min (SD mice) or 6mL/min (CDAHFD mice). The liver was then removed, cut in small pieces and incubated in 25 ml of digestion buffer, complemented with 250µL DNAse I (Roche, 10 104 159 001), at 37°C for 20 min (SD mice) or 25 min (CDAHFD mice). The digested liver suspension was filtered through a 70 µm cell strainer (Falcon™ 352350) and centrifuged at 600g at 4°C for 15 min. Cells were then washed with 50 mL PBS complemented with 120µL DNAse I, centrifuged at 600g at 4°C for 15 min and resuspended in 32 mL of Gey’s Balanced Salt Solution (GBSS/B, 0.4 mM Na_2_HPO_4_, 2.7 mM NaHCO_3_, 0.2 mM KH_2_PO_4_, 137 mM NaCl, 5 mM KCl, 1 mM MgCl2, 0.3 mM MgSO_4_, 5.5 mM Glucose, 2 mM CaCl_2_, pH 7.4) complemented with 120µL DNAse I. Antibiotics were added to buffers (except perfusion buffer) of all steps of cell preparation from CDAHFD-mouse liver (dissociation, gradient step, sorting and immunolabeling): penicillin (200 U/mL), streptomycin (200 µg/mL) (Gibco, 15140-122) and gentamicin (70 µg/mL) (Gibco, 15710-049).

#### HSC isolation using a density gradient

Cells suspended in GBSS/B were mixed with 16 mL Nycodenz solution (Serumwerk, 18003; 0,3 g/L in GBSS/A) and then divided into 4 x 15 mL-tubes, each containing 12 mL of the cell / Nycodenz mixture. GBSS/A buffer has the same composition as GBSS/B buffer without addition of NaCl. A volume of 1.5 mL of GBSS/B buffer was gently added on top of the cell/Nycodenz mixture, using a syringe connected to 26G needle. The tubes were centrifuged at 1380g at 4°C for 17 minutes without brake. HSCs were collected at the interface formed between GBSS/B and GBSS/A washed in PBS, centrifuged at 600 g for 15 min at 4°C and resuspended in 500µL PBS supplemented with 1% FBS (Fetal Bovine Serum). Cells were counted using a Malassez cell.

#### HSC sorting using FACS

The cell concentration was adjusted to 10^7^ cells/mL in PBS-SVF 1% and cell sorting was performed with the MoFlo Astrios (Beckman Coulter, France) at 25 PSI with a 100 µm nozzle at approximately 4000 events per sec. The FSC and SSC were read logarithmically with the 488 nm laser and the threshold has been set on the FSC. For hypHSC sorting, green fluorescence was read with the 488 laser (513/26 band pass classically used for FITC or GFP) and for qHSC sorting, blue fluorescence was read with the 405 nm laser (448/59 band pass classically used for BV 421). The sorting gate of hypHSC was defined on SSC-561 on linear scale vs green fluorescence (488 laser (513/26 band pass) plot for better definition.

In our experience, the number of sorted events given by sorter was always greater than the number of cells, meaning that this number corresponded rather to the number of elements detected by the device than the number of actual cells. Thus, a first sorting of 100 000 events was carried out, to count the number of cells obtained using a Mallassez cell. A % of events corresponding to cells was then obtained and made it possible to determine the number of events to be collected to actually have the number of cells required. For example, if cell sorter indicated that 100 000 events were sorted, and we counted 40 000 cells, we assumed that 40% of the events detected by the sorter corresponded to actual cells. NB: We observed that a % greater than 60% of events corresponding to cells gave a good quality cell preparation.

For qPCR analysis, between 100 000 and 300 000 sorted HSC were suspended in RNA later (Invitrogen, AM7020), flash-frozen in dry ice and stored at -80°C. For immunofluorescent labeling experiments, sorted HSCs were suspended in PBS-SVF 10% or DMEM containing 10 % SVF.

For RNA-seq analysis, 20 000 sorted HSCs were harvested directly from the cell sorter in lysis buffer (Buffer RL #51800, Norgen Biotek Corp.), flash-frozen in dry ice and stored at -80°C.

### 3- Microscopic analysis of cells after gradient or after sorting

HSC quantification was performed by counting fluorescent cells on images acquired with an emCCD camera Retiga 2000 (Photometrics) mounted on an Axiovert (Zeiss, Germany) microscope and analyzed using ImageJ software. Filter set configuration was as follows: blue fluorescence for qHSCs from SD-mouse liver: excitation: 365/12 nm and emission long pass starting at 397 nm; green fluorescence for hypHSCs from CDAHFD-mouse liver excitation: 470/40 nm and emission: 540/50nm. Objectives used were 32X/NA 0,4 and 20X/NA 0,3 (Zeiss, Germany).

The number of total cells was counted under phase contrast illumination and the number of HSCs was determined by counting cells containing droplets emitting blue-only (qHSC) or blue and green (hypHSC) fluorescence on at least 5 microscope’s fields corresponding to 100-200 cells for each condition. For sorted hypHSCs, the number of intact hypHSCs was determined by counting green fluorescent cells with fluorescence contained in intact cytoplasmic droplets, and the number of opaque hypHSCs was determined by counting the number of cells showing fluorescence throughout the cell cytoplasm.

#### Immunostaining of sorted cells

Sorted HSCs were resuspended in DMEM medium (10% SVF) and seeded in microwells mounted on a polymer coverslip (Ibidi) and covered with collagen gel (Collagen type I, 1.5 mg/mL final in NaOH 6.7mM) at a density ranging from 80000 to 120000 cells/cm². Cells were incubated at 37°C and 5% CO2 for 1-2 days, to enable their adhesion to the collagen gel and avoid cell loss during the next steps. The cells were then fixed with 4% para-formaldehyde for 30 minutes at room temperature (RT), rinsed with 1% PBS-BSA, permeabilized (only in the case of intracellular protein labeling) with 0.1 % PBS-Triton X100 for 10 minutes at RT. After rinsing with 1% PBS-BSA, cells were incubated for at least 1 hour in 1% PBS-BSA and then incubated with primary antibodies diluted in 1% PBS-BSA and 0.05%Triton X100 overnight at 4°C. After rinsing with 1% PBS-BSA, cells were incubated with secondary antibodies and DAPI (Sigma-Aldrich, D1388, 1/1000) diluted in 1% PBS-BSA and 0.05% Triton X100 for 1 hour at RT. Primary antibodies were used at the following concentrations: cRBP1 (abcam ab154881; 1/50), anti-Desmin conjugated to AlexaFluor647 (Santa Cruz Biotechnology sc-23879; 1/10); LRAT (ab166784; 1/20), aSMA (A5228; 1/10) and secondary antibody Goat anti-rabbit IgG conjugated to Alexa Fluor 647 (Invitrogen (A-21244); 1/200). Cells were then rinsed with 1% PBS-BSA and mounted in mounting medium (Fluoromount-G; Invitrogen®) up to 24 hours at 4°C. Microscopy acquisitions were performed with laser scanning fluorescent microscope SP8 (Leica Microsystems, Germany) using a 40x/NA 1,3 oil immersion objective. Images were acquired using the software Leica Acquisition System X (Leica Microsystems, Germany) and treated using open source FiJi software.

### 4- Analysis of cells after gradient using flow cytometry

Cells recovered after the gradient were suspended in PBS-SVF10% at a density of 30000 to 50000 cells/100µl and fixed with 2% PBS-PFA for 15 min at RT. After washing by adding 200µL PBS-SVF 10% and centrifugation at 600g at 4°C for 15 min, they were resuspended at the same density in 10 %PBS-SVF and primary and secondary antibodies were added. If the antibody targets an intracellular protein, cells were previously permeabilized by adding 0.05% Triton X100. Tubes were gently shaken by rotation for 40 min at RT. Cells were then washed by adding 200µl of 10% PBS-SVF, centrifuged at 600g at 4°C for 15 min and were resuspended in 10% PBS-SVF at an average of 30000 cells/ 100µL for flow cytometry analysis. The following antibodies were used: at 1/50 dilution, anti-ASGPR 1/2 (FL-291; (sc-28977), anti-RBP1 (ab154881) and anti-Desmin (ab15200), and at 1/200 dilution, anti F4/80 (CI:A3-1) (sc-59171) and anti-CD31 (MEC 13.3) (BD Pharmingen 550274). The following secondary antibodies were used: Goat anti-rabbit IgG, Alexa Fluor 488, (Invitrogen A-11008) at 1/200 and Goat anti-rat IgG, Alexa Fluor 488, (Invitrogen; A-11006) at 1/400. Cell suspensions were analyzed on a flow cytometer Guava Easycyte™ cytometer (Millipore) equipped with two lasers (excitation 488 nm and 642 nm). Unstained cells recovered after gradient were analyzed on a LSRII BD FACS Fortessa flow cytometer (BD Biosciences, Cybio facility, Cochin Institute) equipped with 4 lasers (excitation 355nm, 405 nm, 488 nm and 640 nm). Data were processed using FlowJo software (version 10, Tree Star Inc.).

### 5- Transcriptomic analysis Bulk RNAseq

RNA was extracted from sorted qHSC or hypHSC using the Single Cell RNA purification (kit #51800, Norgen Biotek Corporation, Thorold, ON, Canada) according to the manufacturer’s protocol and eluted in 23µL. After extraction, RNA concentrations were obtained using fluorometric Qubit RNA assay (Life Technologies, Grand Island, New York, USA). The quality of the RNA (RNA integrity number) was determined on the Agilent 2100 Bioanalyzer (Agilent Technologies, Palo Alto, CA, USA) as per the manufacturer’s instructions using pico chip kit.

To construct the libraries, 2.5 ng of high-quality total RNA sample (RIN >7) was processed using NEBNext low input RNA library prep kit (NEB) according to manufacturer instructions. Briefly, total RNA molecules are reverse-transcribed using oligo dT primers and template switching oligos. cDNA was then amplified with 3’ and 5’ adapters and QC controlled. After fragmentation, amplified cDNA is then end repaired and ligated with NEBNEXT adapters. PCR enrichment with single barcode was then realized to obtain the final library. Libraries were then quantified with Qubit HS DNA assay (Life Technologies, Grand Island, New York, USA) and library profiles assessed using the DNA High Sensitivity LabChip kit on an Agilent 2100 Bioanalyzer (Agilent Technologies, Palo Alto, CA, USA). Libraries were sequenced on an Illumina Nextseq 2000 instrument using 59 base-lengths read V2 chemistry in a paired-end mode.

RNA-Seq data analysis was performed by GenoSplice technology (www.genosplice.com). Analysis of sequencing data quality, reads repartition (*e.g.*, for potential ribosomal contamination), inner distance size estimation, genebody coverage, strand-specificity of library were performed using FastQC v0.11.2, Picard-Tools v1.119, Samtools v1.0, and RSeQC v2.3.9. Reads were mapped using STAR v2.7.5a [PMID: 23104886] on the Mouse mm39 genome assembly and read count was performed using featureCount from SubRead v1.5.0 and the Mouse FAST DB v2022_1 annotations. Gene expression was estimated as described previously [PMID: 34495298]. Only genes expressed in at least one of the two compared conditions were analyzed further. Genes were considered as expressed if their FPKM value was greater than FPKM of 98% of the intergenic regions (background). Analysis at the gene level was performed using DESeq2 [PMID: 25516281]. Genes were considered differentially expressed for log2FC ≥ 1 and adjusted p-values ≤ 0.05. Overrepresented analyses were performed using WebGestalt v0.4.4 [PMID: 31114916] merging results from up-regulated and down-regulated genes only, as well as all regulated genes. Pathways and ontologies were considered significant with p-values ≤ 0.05.

#### qPCR

Cells sorted and recovered in RNA later were centrifuged at 5,000g at 4°C for 10min. RNA was then extracted using the RNeasy kit (Qiagen 741004) according to provider’s recommendations and eluted in 30 µL of RNAse-free water. Extracted RNA was first treated with DNAse I (Takara, 2270A) to remove genomic DNA and complete removing was controlled by PCR. RNA concentration was then determined using the Quant-iT RiboGreen RNA kit and reagent (Invitrogen, R11490). 10 pg to 5 µg of RNA were mixed with 1 µL of 50 µM random primers, 1 µL of 10 mM dNTP and RNAse-free water to give a final volume of 13 µL and incubated at 65°C for 5 minutes to allow primer hybridization, then on ice for 1 minute. 4 µL of SSIV Buffer, 1 µL of 100 mM DDT, 1 µL of RNAse OUT (Invitrogen, 10777019) and 1 µL of Reverse Transcriptase SuperScript IV enzyme (SSIV, Invitrogen, 18090010) were added to the mixture, which was then incubated at 23°C for 10 minutes, then at 55°C for 10 minutes to allow cDNA synthesis, and finally at 80°C for 10 minutes to inactivate the enzyme. cDNAs were stored at -80°C. For qPCR analysis, 1 µL of diluted cDNA (1:10 in RNAse-free water) was deposited in 384-well microplate to which is added 19 µL of a mixture containing: 10 µL of Power SYBER Green Master Mix (Thermo Fisher Scientific, 10219284), 0.1 µL of the sense strand primer 0.1 µL of the anti-sense strand primer (See Table S2) and 8.8 µL of RNAse-free water. qPCR was performed on the BIO-Rad CFX 384, according to the following steps: enzyme activation by heating the plate at 95°C for 10 minutes, followed by 40 cycles of denaturation at 95°C for 15 seconds and hybridization and elongation at 60°C for 1 minute. Results were analyzed using Bio-Rad CFX Maestro software. The stability of housekeeping genes was assessed on samples of sorted hypHSCs and qHSCs, using the GeNorm algorithm of the “Reference Gene Selection” tool in Biorad’s CFX software. This analysis validated 18S, HPRT1, Actin and PGK1 as the 4 reference genes for studying the expression of genes of interest in different HSC populations.

## Supporting information

Heckmann_supp_informations

## Acknowledgments

We thank Servane Le Plénier and her team from Animal facility - UAR 3612 CNRS - US25 Inserm – Faculté de Pharmacie de Paris Cité, Université de Paris Cité, Paris, France, for technical support in animal experiments. We thank the Plateforme d’Imagerie Cellulaire et Moléculaire PICMO, US25 Inserm, UAR3612 CNRS, Université Paris Cité, Faculté de Pharmacie, Paris, France. We thank the Plateforme de cytométrie et d’immunologie CYBIO, Université Paris-Cité for preliminary sorting experiments. We deeply thank Pierre-Henri Commere from cytometry Platform, Institut Pasteur, Université Paris Cité, Paris, France for strong help in cell sorting. We thank the Genomics Core Facility of the Institute Cochin for NGS data production and analysis. We wish to thank Dr. Pierre de la Grange (GenoSplice, Paris) for RNA-seq analyses and bioinformatics support. We gratefully acknowledge the UTechS Photonic BioImaging (Imagopole), C2RT, Institut Pasteur, supported by the French National Research Agency (France BioImaging, ANR-10-INBS-04; Investments for the Future).

## Authors contributions

M.H., NEH. D. and C. H. performed the experiments and analyzed the data; PH. C provided resources for FACS, advices and performed the sorting; J. F. provided microscopy resources, advices and performed microscopy acquisitions; G. S. performed experiments and analysis on mice microbiota; M.H., P. B., V.E and C.H. performed formal analysis on RNAseq; V.E. and C.H. designed the methodology, supervised the project and wrote the manuscript with input from all authors.

## Data Availability Statement

The RNA-sequencing data have been deposited in GEO database under the accession number GSE294603. All other relevant data are within the paper and its Supplemental Information files.

