## Supplementary material for "Purification protocol of hypertrophied hepatic stellate cells for their transcriptomic characterization from CDAHFD mice liver": Heckmann_supp_informations

##### Supplemental Methods

##### Fecal and hepatic sample collection and genomic DNA extraction

Mouse fecal pellets were collected at 0, 3, 6, 9 and 12 weeks of diet from CDAHFD fed mice (n=4 to 8) and SD fed mice (n=5) flash-frozen and stored at -80°C. Liver frozen samples were lysed for 2 h in solution containing 25 µL of proteinase K (Qiagen) and 180 µL of ATL lysis buffer (Qiagen). Genomic DNA (gDNA) from fecal samples and liver tissue lysates were extracted according to the recommendations of the International Human Microbiome Standards (IHMS: http://www.human-microbiome.org/). Briefly, aliquot of each sample was suspended in 250 µl of guanidine thiocyanate, 40 µl of 10% N-lauroyl sarcosine, and 500 µl of 5% N-lauroyl sarcosine. Genomic DNA was extracted by mechanical disruption of the microbial cells with glass beads (0.1 mm, Sigma-Aldrich) using a TissueLyser III (QIAGEN), and nucleic acids were recovered from clear lysates by alcohol precipitation as previously described[1]. The concentration and purity of each gDNA sample obtained are assessed using a nanodrop (Nanodrop One, Thermo Fisher Scientific) and the quality of the DNA is assessed by means of a genomic gel (1% agarose, 100 volts, 1 hour) as recommended by the IHMS. Any sample showing poor purity or excessive degradation is excluded from the study.

##### Microbial load analysis

Extracted genomic DNA was used to amplify the V4 region of the 16S rRNA gene by standard and quantitative real-time PCR, using universal primers for counting the microbial load. Universal primers located in highly conserved sequences targeting the V4 hypervariable region of the 16S rRNA genes (290 bp) were used: V4F_517_17 (5'-GCC AGC CGC GGT AA-3') and V4R_ 805_19 (5'-GAC TAC CAG GGT ATC TAA T-3').

Standard PCR (0.15 units of Taq polymerase (AmpliTaq Gold, Life Technologies®) and 20 pmol/µL of the forward and reverse primers (IDT Technologies®) was run in a Mastercycler gradient (Eppendorf) at 94°C for 3 min, followed by 35 cycles of 94°C for 45s, 56°C for 60s, 72°C for 90s and a final cycle of 72°C for 10 min.

In order to assess the microbial load, the extracted DNA was used to amplify the V4 region of the 16S rRNA gene by quantitative real-time PCR (qPCR) using universal primers for counting microbial load as previously described[2]. The PCR was performed in a volume of 25 µl using Power SYBR green PCR master mix (Fisher Scientific) containing 100 nM (each) primer. The reaction conditions were 50°C for 2 min, 95°C for 10 min, and 40 cycles of 95°C for 15 s and 60°C for 1 min. All samples were analyzed in triplicate and mean values were calculated. Reading and analysis of results was evaluated using Bio-Rad CFX Manager 3.1 software (Bio-Rad).

### Supplemental Figures – Figure S1

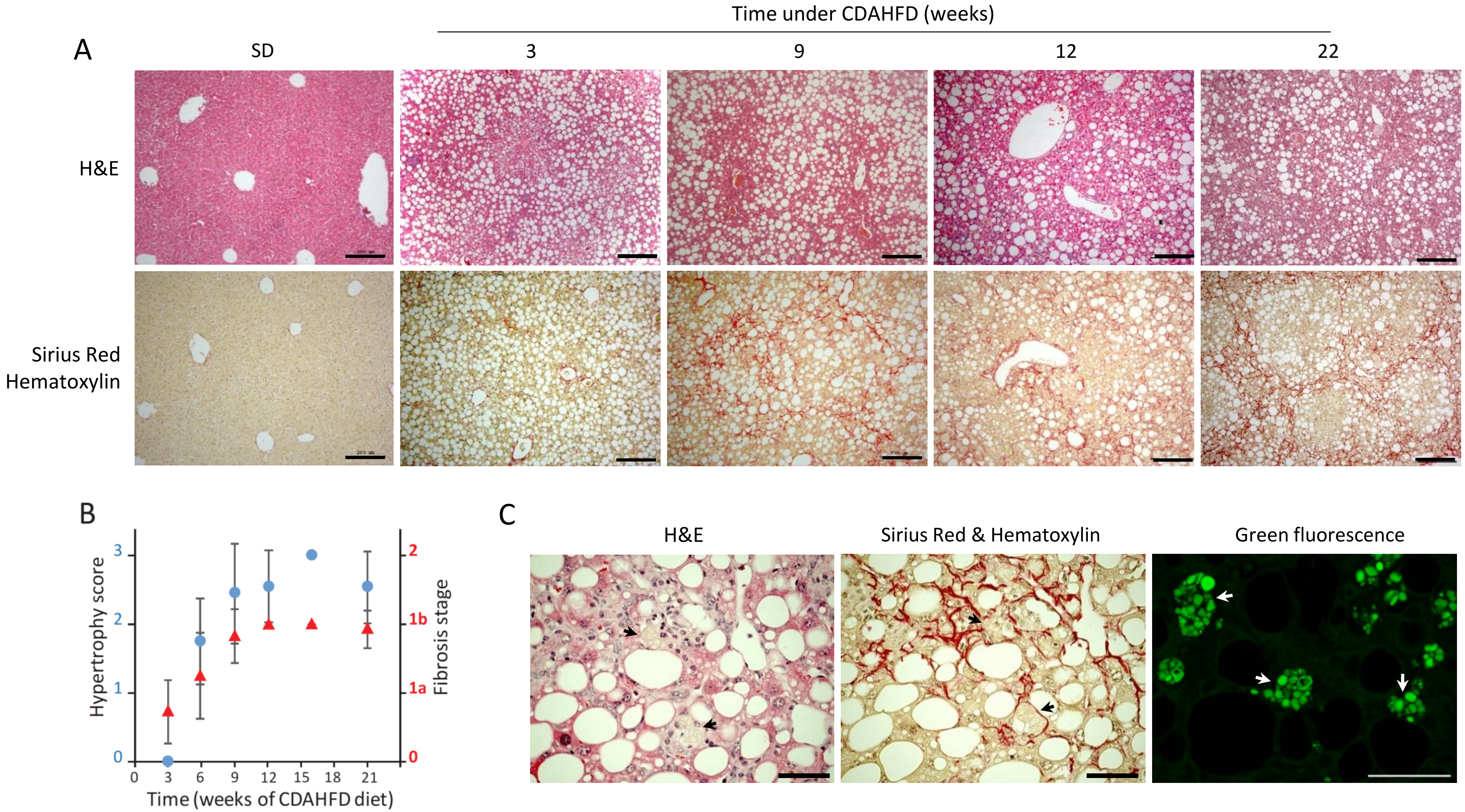


**Figure S1: Kinetic of fibrosis and HSC hypertrophy progression as a function of CDAHFD time.** A) Histological colorations of CDAHFD and SD-fed mice paraffin liver slices (top: Hematoxylin/Eosin, low: Sirius Red). B) HSC hypertrophy (blue dots) and fibrosis (red triangles) scores as a function of CDAHFD time. (n = 10–14 for each time point, means ± standard deviation). (data from Hoffmann *et al.*, Scientific Reports, 2020) C) Examples of HSC hypertrophy (pointed by an arrow) on CDAHFD-mouse liver observable on histological sections stained with H&E, Sirius Red and on unstained paraffin slices by fluorescence microscopy (λexc= 488 nm, λem= 505-535 nm, green). Scale bar : 50µm

### Supplemental Figures – Figure S2

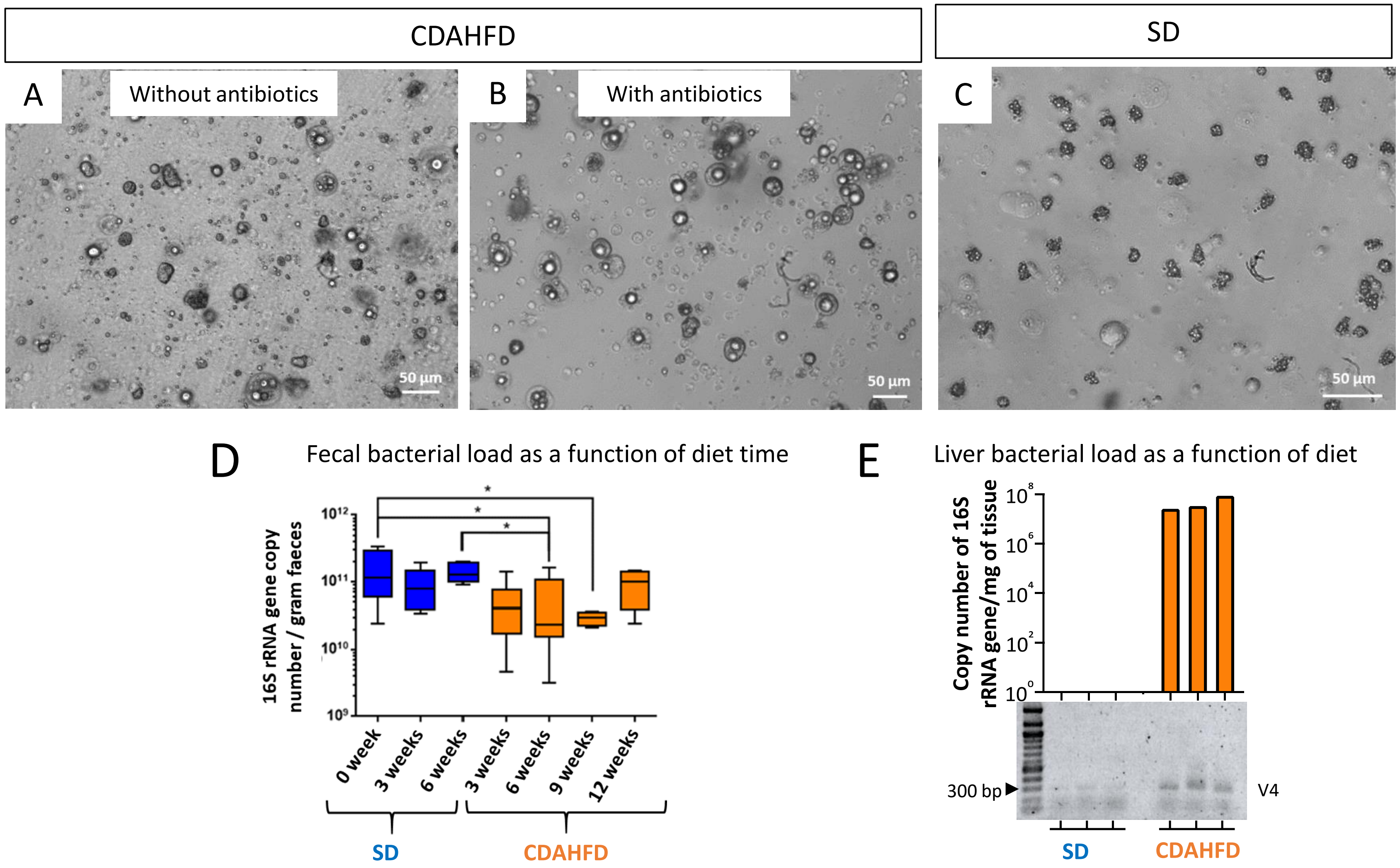


**Figure S2: Bacterial contamination of cell preparations obtained from CDAHFD mice. A-B) Cell preparations from CDAHFD mouse without (A) and with (B) antibiotics** during HSC purification. Cell preparation with antibiotics contains less bacteria than cell preparation without antibiotics in buffers. **C) Example of cell preparation from SD mouse** without antibiotic. **D) Evaluation of fecal bacterial load as a function of diet time.** Representation of the number of copies of the gene encoding 16S rRNA obtained after amplification by qPCR. Statistics: n = 5 for each SD diet time, n = 8 for 3, 6 and 9 weeks of CDAHF diet and n = 4 for 12 weeks of CDAHF diet, mean ± SD, Mann-Whitney test, *p<0.05. **E) Evaluation of liver bacterial load as a function of diet.** Graph representative of the number of copies of the gene encoding 16S rRNA obtained after amplification by qPCR for 3 liver samples from SD-fed mice and 3 CDAHFD-fed mice (top). Image of genomic gel of corresponding samples after amplification of V4 region of the 16S rRNA gene by standard PCR (bottom).

### Supplemental Figures – Figure S3

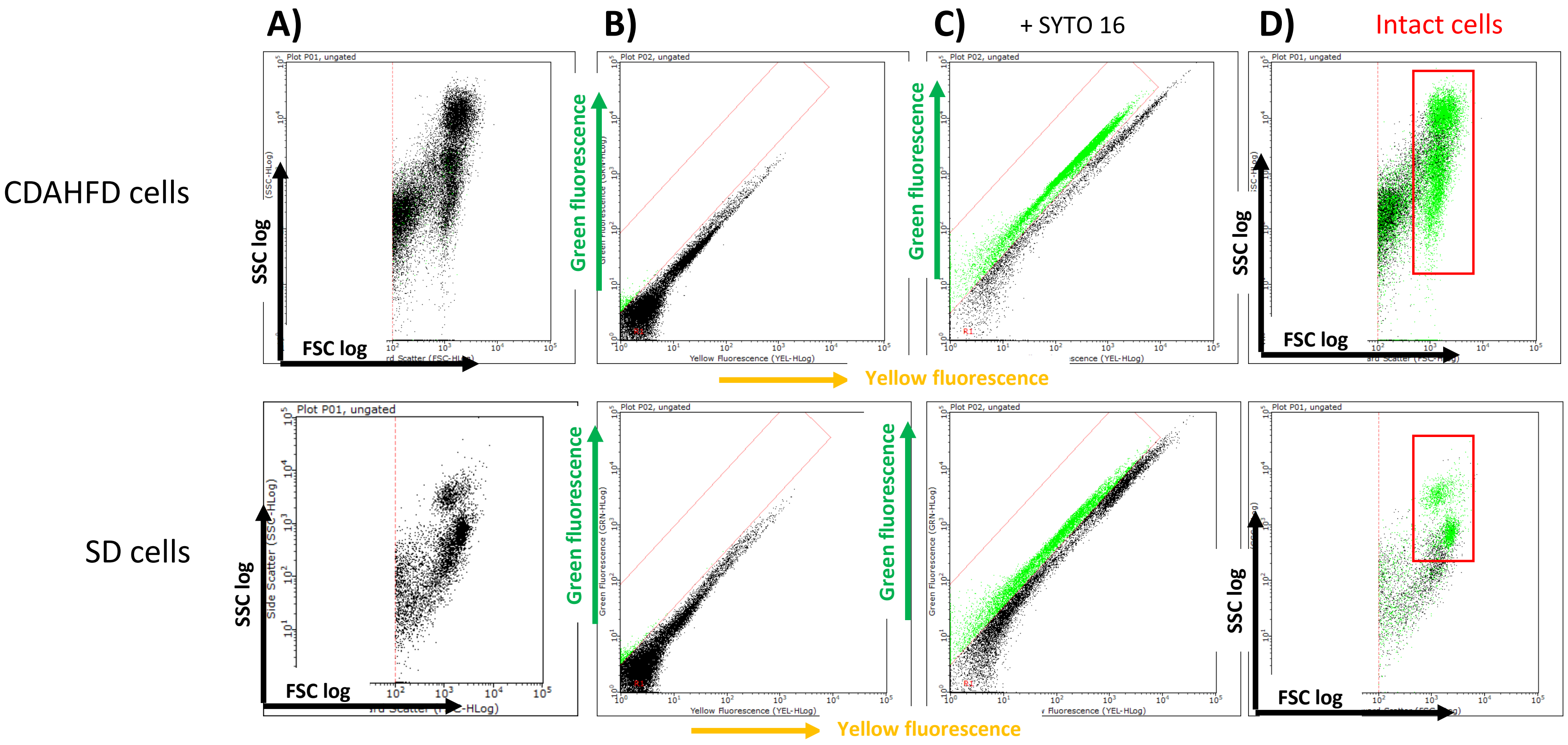


**Figure S3 : Localization of cells obtained from CDAHFD- and SD-mouse liver after density gradient on FSC/SSC dot plot.** A) Dot plot of cell preparations analyzed as a function of FSC and SSC. B) Dot plot of the same cell preparations without labeling, analyzed according to green fluorescence (λexc = 488 nm and λem = 525 nm) and yellow fluorescence (λexc = 488 nm and λem = 583 nm). C) Dot plot of the cell preparations labeled with SYTO16. The points corresponding to the positive SYTO16 labelling are represented in green. D) The points corresponding to the syto16-labelled events, therefore to intact cells are represented in green on the FSC-SSC dot plot.

### Supplemental Figures – Figure S4

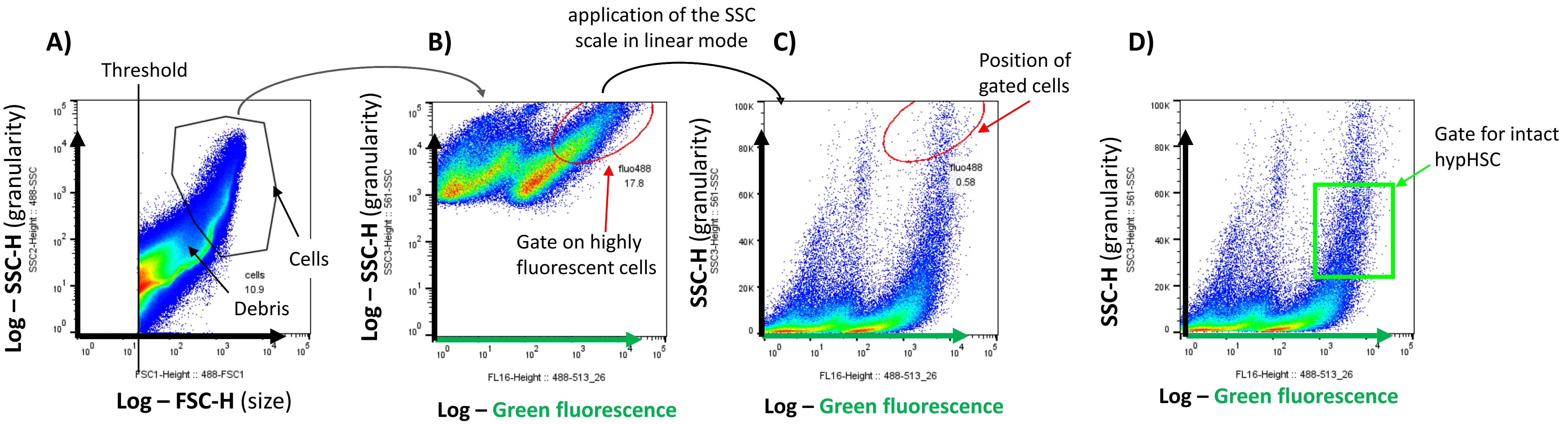


**Figure S4. SSC scale adjustment strategy for a precise selection of hypHSC on SSC/green fluorescence dot plot.**
(A) Dot plot log-SSC/log-FSC allowing to determine the gate for cells (B) dot plot for SSC (log) /green fluorescence (log) and gating strategy (red circle) to highly green fluorescent cells to hypHSC C) application of the SSC scale in linear mode on dot plot for SSC/log-green fluorescence showing position of selected cells on the top of the dot-plot of highly fluorescent cells D) Representation of the ideal gate for hypHSC recovering on dot-plot SSC (lin)/green fluorescence (log)

### Supplemental Figures – Figure S5

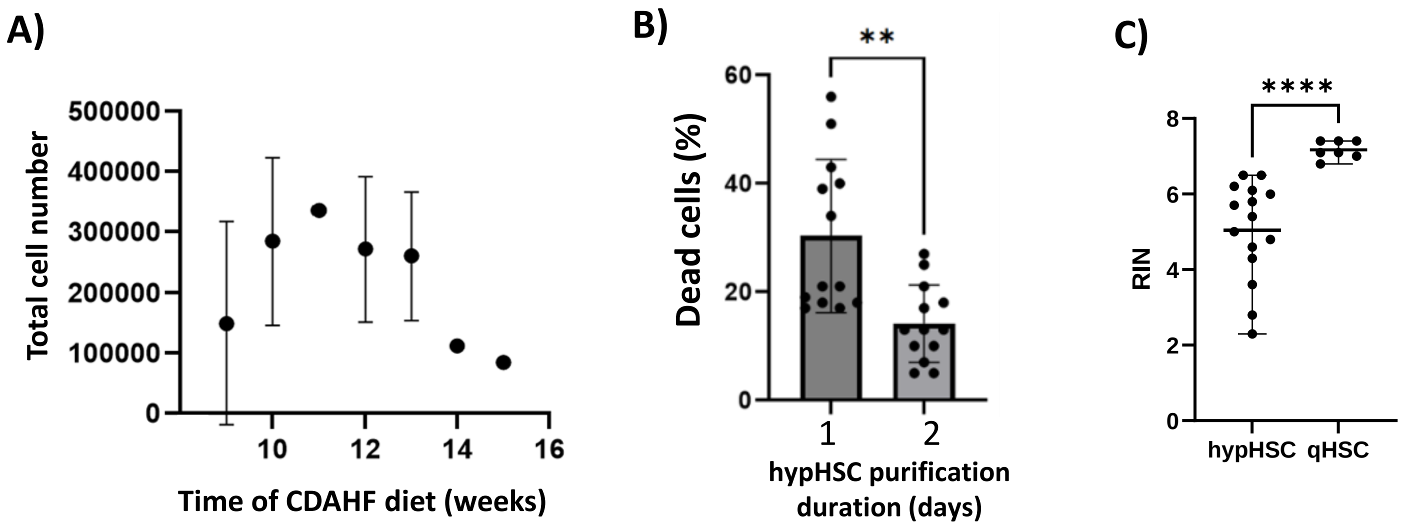


**Figure S5 : A) Number of recovered cells after FACS as a function of CDAHFD time.** Statistics: for 11 weeks n=1; for 9, 14 and 15 weeks : n=2, for 10 and 13 weeks and for 12 weeks: n=8, mean ± standard deviation, Kruskall-Wallis test not significant. **B) Mortality of hypHSCs collected after sorting according to the duration of purification** (1 or 2 days). Statistics: n = 13 CDAHFD mice for both conditions, mean ± SD, **p<0.005, Mann-Whitney test. **C) RIN values of RNAs extracted from sorted hypHSC and qHSC for RNAseq.** hypHSC n=15; qHSC n=7. ****p<0.0001 Mann-Whitney test

### Supplemental Figures – Figure S6

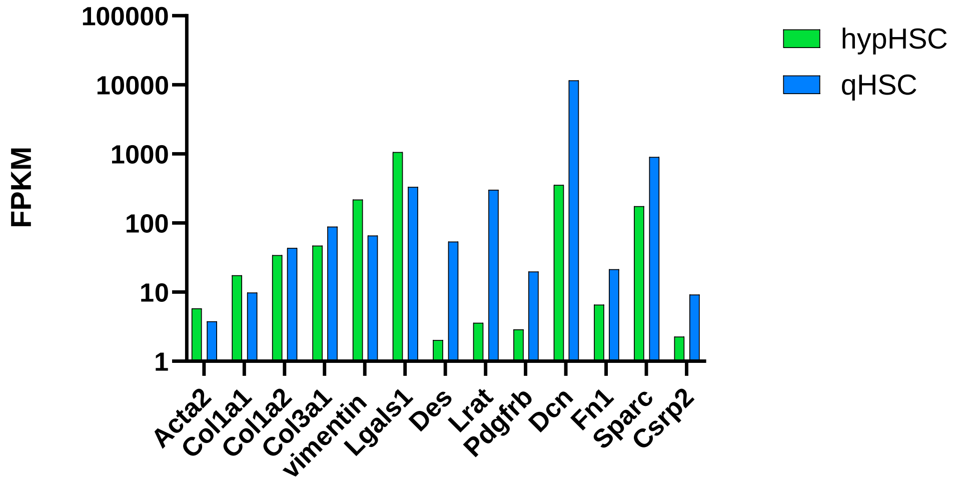


**Figure S6. Analysis of the expression of some HSC marker genes by quiescent and hypertrophied HSCs.** FPKM (Fragments Per Kilobase of transcription per Million mapped reads) mean of 5 samples for each condition. Acta2= α-SMA; Col1a1, Col1a2 and Col3a1 = collagens 1a1, 1a2 and 3a1; Vim=vimentin; Lgals1= galectin-1; Des= desmin; Lrat= lecithin retinol acyltransferase*;* PDGFRb= PDGFR-β; DCN = decorin; Fn1 = Fibronectin ; Sparc = osteonectin Csrp2 = cysteine- and glycine-rich protein.

### Supplemental Figures – Figure S7

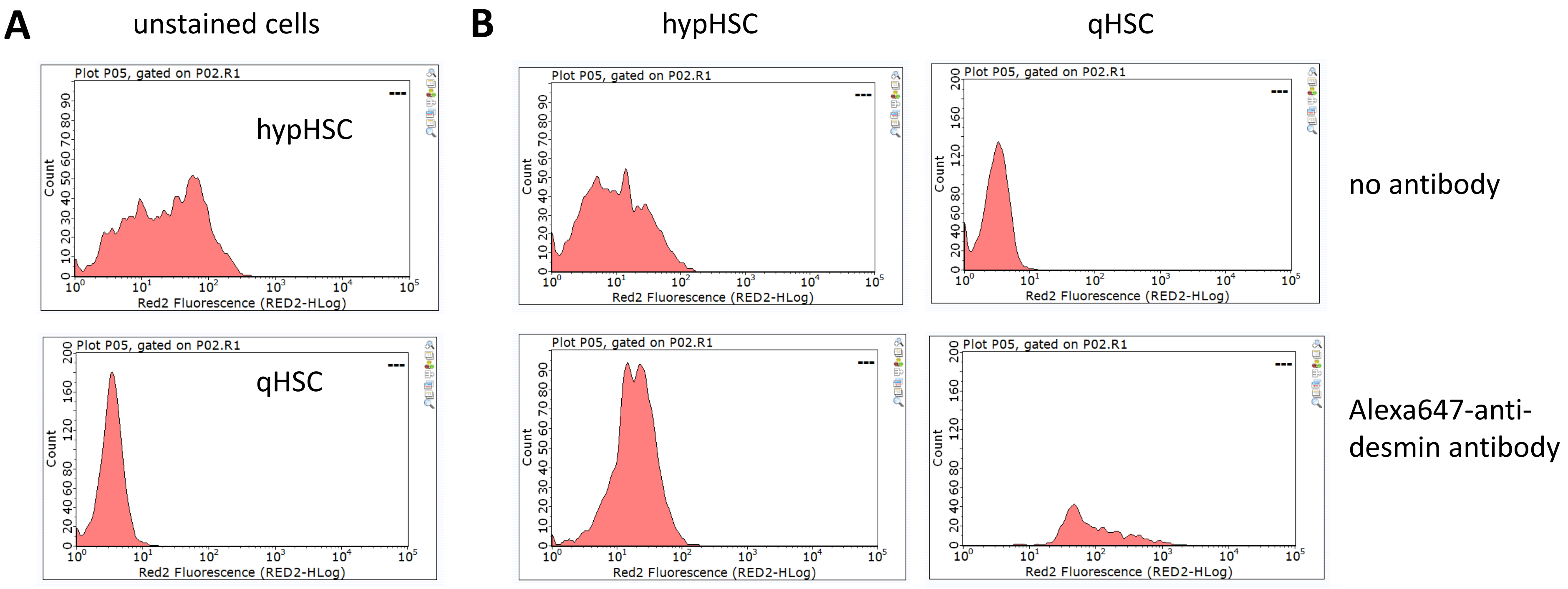


**Figure S7. Representative histograms from flow cytometry analysis of sorted hypertrophied and quiescent HSC.** Hypertrophied (hypHSC) and quiescent (qHSC) hepatic stellate cells were analyzed by flow cytometry and Red2 fluorescence signals were recorded and displayed as histograms. A) unstained cells. B) unstained cells (up) and cells stained with Alexa647-labeled anti-desmin antibody (down).

HypHSC show a very broad fluorescence signal, much more wide-ranging than that of qHSCs. When these two cell types are analyzed by flow cytometry, e.g. with excitation at 647 nm, qHSCs show a classic peak in the low range of fluorescence intensity, whereas hypHSCs show a much wider distribution of cells as a function of fluorescence intensity, with a heterogeneous profile. When both cell types are incubated with an anti-desmin antibody coupled to the Alexa 647 fluorophore, the qHSC peak shifts towards higher intensities, univocally demonstrating labeling of all cells. For hypHSCs, the shape of the distribution profile changes, but the range of intensities remains unchanged, making it impossible to demonstrate specific labeling.

### Supplemental Tables

#### Table S1

| gene name | regulation (up/down/NR) | Fold change | Log2(FoldChange) | P-value | hypHSC | | qHSC | |
|---|---|---|---|---|---|---|---|---|
| | | | | | expression | Rank | expression | Rank |
| adipor1 | up | 2,57 | 1,36 | 6,97E-13 | 10,57 | 4033 | 5,18 | 7025 |
| sparc | down | 3,94 | -1,98 | 1,53E-04 | 175,7 | 440 | 900,2 | 58 |
| dcn | down | 25,27 | -4,66 | 4,65E-65 | 357,15 | 209 | 11620,53 | 2 |
| csrp2 | down | 3,14 | -1,65 | 1,16E-05 | 2,27 | 9045 | 9,25 | 5127 |
| bambi | down | 2,67 | -1,42 | 8,75E-04 | 0,78 | 13334 | 2,74 | 9305 |
| plin2 | up | 3,43 | 1,78 | 4,77E-10 | 733,77 | 93 | 270,2 | 252 |
| vim | up | 4,33 | 2,11 | 4,52E-15 | 218,77 | 338 | 65,76 | 1175 |
| pparg | up | 90,29 | 6,50 | 1,30E-13 | 11,66 | 3793 | 0,17 | 23446 |
| lrat | down | 66,45 | -6,05 | 6,44E-53 | 3,59 | 7345 | 301,73 | 217 |
| gfap | down | 43,48 | -5,44 | 1,73E-09 | 0,27 | 19337 | 14,17 | 4013 |
| des | down | 20,82 | -4,38 | 5,58E-14 | 2,03 | 9500 | 53,68 | 1421 |
| acta2 | NR | | | | 5,81 | 8071 | 3,78 | 5725 |
| col1a1 | NR | | | | 17,54 | 2882 | 9,86 | 4963 |
| col1a2 | NR | | | | 34,41 | 1795 | 43,67 | 1690 |
| col3a1 | NR | | | | 46,92 | 1410 | 88,44 | 866 |
| notch3 | NR | | | | 0,07 | 25761 | 0,13 | 24490 |
| timp1 | NR | | | | 50,73 | 1327 | 30,77 | 2267 |

**Table S1. Regulation of HSC specific genes.** Table of HSC specific genes with regulation in hypHSC compared to qHSC, value of fold-change, p-value and corresponding expression value (FPKM) and rank for hypHSC and qHSC. NR = unregulated, up = up-regulated (pink), down = down-regulated (green).

### Supplemental Tables

### Table S2

| Gene | Sense strand | Antisense stand |
|---|---|---|
| **Fn1** | GAGAGGAGTGGGAGCGGTTG | TCCCTTTCCATTCCCGAGGC |
| **Laminin** | GCCGGGTGAGGAGAACAAAGTA | TGGAGAGGTAGCGGGGAGAAA |
| **Pdgfra** | AAACAAACGGAGGAGCTGCG | CCCCATAGCTCCTGAGACCTTC |
| **Pdgfrb** | ACGTGGACCCTGTGCAGTTG | GAGTGCGTCCCAGAACAAGC |
| **Itgav** | CTTTGGGCTGTGGAATCGCC | AGGATTGCGCTCTTGCCTCT |
| **IL1b** | TCACAAGCAGAGCACAAGCC | GCATTAGAAACAGTCCAGCCC |
| **18S** | TTGACGGAAGGGCACCACCAG | GCACCACCACCCACGGAATCG |
| **PGK1** | GCTGAACTCAAATCTCTGCTG | TCTTTTCCCTTCCCTTCTTCC |
| **HPRT1** | GCTTACCTCACTGCTTTCC | TTCATCATCGCTAATCACGAC |
| **MMP12** | GATGAGGCAGAAACGTGGAC | TGGGGTACATTATTGACTTTGGA |
| **LRAT** | GTCCACAGGCTGAGAAGTTCTAT | AATCCCAAGACAGCCGAAGCA |
| **Col1a1** | GTC GCT TCA CCT ACA GCA C | CAA TGT CCA AGG GAG CCA C |
| **Col5a1** | CAA AGG TGA AAA GGG CCA TC | CTC CTT TAG GAC CAG ATG AAC C |
| **Timp 1** | TCCTAGAGACACACCAGAGCAGA | GGGGAACCCATGAATTTAGCCC |
| **Timp 2** | GGCTGTGAGTGCAAGATCACT | CACGCGCAAGAACCATCACTT |
| **aSMA** | GGCTCTGGGCTCTGTAAGG | CTCTTGCTCTGGGCTTCATC |
| **Vegfa** | TTACTGCTGTACCTCCACC | ACAGGACGGCTTGAAGATG |
| **TNF** | AGAAAAGCAAGCAGCCAACC | CATAGGCACCGCCTGGAGT |
| **Thsb2** | CATCGGTGCCAAGCAGTTCC | ACTTGCGGTCCTGCTTCAGT |
| **Actine** | GCTTCTTTGCAGCTCCTTCG | ACCCATTCCCACCATCACAC |

**Table S2: Primer sequences used for qPCR analysis of gene expression**
